# Grapevine cell response to carbon deficiency requires transcriptome and methylome reprogramming

**DOI:** 10.1101/2024.02.02.578586

**Authors:** Margot M J Berger, Virginie Garcia, Nathalie Lacrampe, Bernadette Rubio, Guillaume Decros, Pierre Pétriacq, Amélie Flandin, Cédric Cassan, Ghislaine Hilbert-Masson, Sophie Colombié, Rossitza Atanassova, Philippe Gallusci

## Abstract

Sugar limitation has dramatic consequences on plant cells, which include a profound reorganization of the cell metabolism, transcriptional reprogramming, and the recycling of cellular components to maintain fundamental cell functions. There is so far no description of the possible contribution of epigenetic regulations in the adaptation of plant cells to limited carbon availability. We investigated this question using non-photosynthetic grapevine cells (*Vitis vinifera*, cv Cabernet Sauvignon) cultured *in vitro* with contrasted glucose concentrations. As expected, limited sugar availability in the culture medium led to a rapid cell growth arrest. This was associated with a major metabolic shift characterized by depletion in soluble sugar and total amino acids, an increase in malate content and changes in the cell redox status. Consistently, flux modeling showed a dramatic slowdown of many pathways required for biomass accumulation such as cell wall polymers and total protein content. In contrast, anaplerotic fluxes, the synthesis of some amino acids, redox and polyamine metabolism were enhanced.

Carbon deprivation also resulted in a major transcriptional reprogramming characterized by the induction of genes involved in photosynthesis, and the repression of those related to sucrose mobilization or cell cycle control. Similarly, the epigenetic landscape was deeply modified. Glucose-depleted cells showed a higher global DNA methylation level than those grown with glucose. Changes in DNA methylation mainly occurred at transposable elements, but also at genes including differentially expressed genes, suggesting that DNA methylation could participate in the adaptation of cells to limited sugar availability. In addition, genes encoding histone modifiers were differentially expressed suggesting that additional epigenetic mechanisms may be at work during the response of plant cells to carbon shortage.

## INTRODUCTION

Epigenetics is currently defined as the study of heritable and/or stable changes in gene expression that occur without modifications of the DNA sequence (Eichten, Schmitz, and Springer 2014). Epigenetic regulations play important roles in the development of plants, as well as in their response and adaptation to changing environments or stresses (H. Zhang, Lang, and Zhu 2018; Lämke and Bäurle 2017). Among the different mechanisms underlying epigenetic regulations, DNA methylation, which is found in most eukaryotic genomes, has been extensively studied during the last decade (Su, Han, and Zhao 2011). DNA methylation is controlled by the balanced activities of DNA-methyltransferases (DNMTs) and DNA-demethylases (DMLs), respectively involved in the addition or the removal of a methyl residue on the C5 of cytosines and in the removal of the methylated cytosine (Law and Jacobsen 2010). In plants, cytosine methylation can occur in the two symmetrical CG and CHG, and in the asymmetrical CHH (H = A, C or T) sequence contexts. Establishment of DNA methylation in all sequence contexts is performed by the RdDM pathway that requires the Domain Rearranged Methyltransferases 1, 2 (DRM1/2), DRD1 and 24nt-long small RNAs, and by the chromomethylase 2 (CMT2) associated with Decrease in DNA Methylation (DDM1) for CHH in constitutive heterochromatic regions (Zemach et al., 2013).

During DNA replication, the newly synthetized DNA strand is not methylated at CG and CHG symmetrical sites, generating hemi methylated DNA molecules. Methylation at hemi methylated CG sites relies on the activity of MET1 together with VIM1, 2 and 3, and at CHG on CMT3. Similar to *de novo* methylation, re-methylation of the newly synthesized DNA strands at non-methylated CHH sites is mediated by both the RdDM pathway and CMT2. CMTs are also dependent on histone methylation mediated by KYP and SUVH5 and 6 through a self-reinforcing loop (Zhang et al., 2018).

It is only recently that potential links between epigenetic regulations and metabolism were evidenced essentially in animal systems (Lu and Thompson 2012). Indeed, epigenetic modifications, both DNA methylation and histone post-translational modifications (HPTMs) are mediated by enzymes, that require metabolic precursors such as the acetyl coenzyme A (acetyl-CoA) or the S-adenosyl-methionine (SAM) and cofactors (NAD+, acetyl-CoA). Epigenetic processes are therefore intimately connected to the metabolism of cells (Shen, Wei, and Zhou 2015; Leung and Gaudin 2020). As far as plants are concerned, impairing the synthesis of metabolic precursors necessary for epigenetic regulations has dramatic consequences on epigenomes (Séré and Martin 2020). For example, mutants affected in organelle or cytoplasmic acetyl-CoA biosynthesis present changes in their epigenetic landscape (Lishuan Wang et al., 2019). Similarly, mutations or pharmacological approaches that impair the 1C metabolism, thereby SAM availability, induce global DNA methylation changes that impact gene expression and plant phenotypes (H. Zhang et al., 2012; Lei Wang et al., 2017). As these metabolic precursors are provided by the central metabolism, its alteration may impact precursor and/or cofactor availability to epigenetic regulations.

It has been described that carbon starvation initiates complex signal transduction processes (Sakr et al. 2018), resulting in significant changes in the plant cell metabolism, resulting in decreased carbohydrate levels, protein degradation, and a global decrease of metabolites and enzymatic activity leading to cell disorganization and death (Morkunas et al., 2012; Sakr et al., 2018; Zhang, Hardie, and Lin 2020).

However, there is no clear description of how the metabolism dynamic is influenced by limitations in carbon availability. This can be achieved by calculating flux analysis as it allows revealing carbon inputs, such as glucose, to condition end-products as key drivers of metabolic behavior (Sweetlove, Obata and Fernie, 2014). The calculation of metabolic fluxes, as the last step to characterize the phenotype, consists of integrating experimental data in constraint-based model to evaluate the dynamics of metabolism and compare the physiological state of cells (Clark et al., 2020). Primary metabolism can be described by medium-scale models reconstructed from biochemical and bibliographic knowledge (Colombié et al., 2015), including specialized or secondary metabolic pathways (Lacrampe et al., *in review*). However, there is no description of the changes in metabolic fluxes induced by carbon shortage.

In addition, cells under carbon shortage undergo important transcriptome reprogramming (Contento, Kim, and Bassham 2004). Genes encoding proteins involved in signal transduction including protein kinases, and transcription factors, but also those coding for enzymes involved in carbon mobilization and membrane transporters were upregulated. In contrast, those encoding proteins involved in translation or cell division were repressed, consistent with an arrest in cell growth and division under carbon starvation (Contento, Kim, and Bassham 2004), ultimately resulting in cell death (Chae and Lee 2001; Azevedo et al., 2014).

Finally, there is growing evidence of an interplay between sugar availability and epigenetic modifications in eukaryotes (Donohoe and Bultman 2012), including plant cells (Chen et al., 2022). For example, in Yeast there an interplay between genome-wide distribution of histone acetylation and transcriptomic changes has been demonstrated following sugar starvation (Hsieh et al., 2022). Similarly, in yellowfin seabream modification of the global DNA methylation level and distribution is considered as a consequence of starvation (Lin et al., 2022). Finally, recent work in Arabidopsis using Rapamicyne, an inhibitor of the Target Of Rapamycin (TOR) kinase, which upon activation by nitrogen and carbon metabolites, promotes energy-consuming processes, affects the methylation distribution at genes involved in carbon and amino acid metabolism, and thereby hormone synthesis, suggesting that the Glc-TOR effect is mediated in part by DNA methylation (Zou et al., 2020).

In the present work, we investigated the possible consequences of carbon deficiency on the methylation landscape of heterotrophic grapevine cells. Cell phenotypes are described by determining cell growth, metabolic profiles and metabolic fluxes. We also described the molecular response of cells based on gene expression analyses and DNA methylation landscape. This multi-omics approach provides pioneer and integrated overview of the response of heterotrophic cells to carbon limitation, from phenotype to DNA methylation, and demonstrated tight interactions between cell primary metabolism, gene expression reprogramming, and DNA-methylation changes. This suggests that epigenetic regulations are central to the response of cells to sugar starvation.

## RESULTS

### Sugar depletion causes a rapid cell growth arrest

Cabernet Sauvignon (CS) cells grown with 110 mM glucose (20 g/L), present a classical latent phase, followed by a growth phase starting at day 3 (D3) and a stationary phase from D8 over (Figure 1A). To evaluate the impact of limited carbon availability, the cells were transferred after 4 days in culture to either a glucose-rich (G+) or a glucose-poor (G-) medium (Figure 1B). The G+ cells displayed a growth phase from D4 to D10 that resulted in an ∼3,5-fold increase in FW, to reach 530.2 (±18.7) g FW/mL at D10 (Figure 1B). In contrast, G-, cell FW showed a 1.2-fold increase, from 160 (±8.93) gFW/mL at D4 to 197.6 (± 9.61) gFW/mL at D5, with no further significant changes.

**Figure 1.**
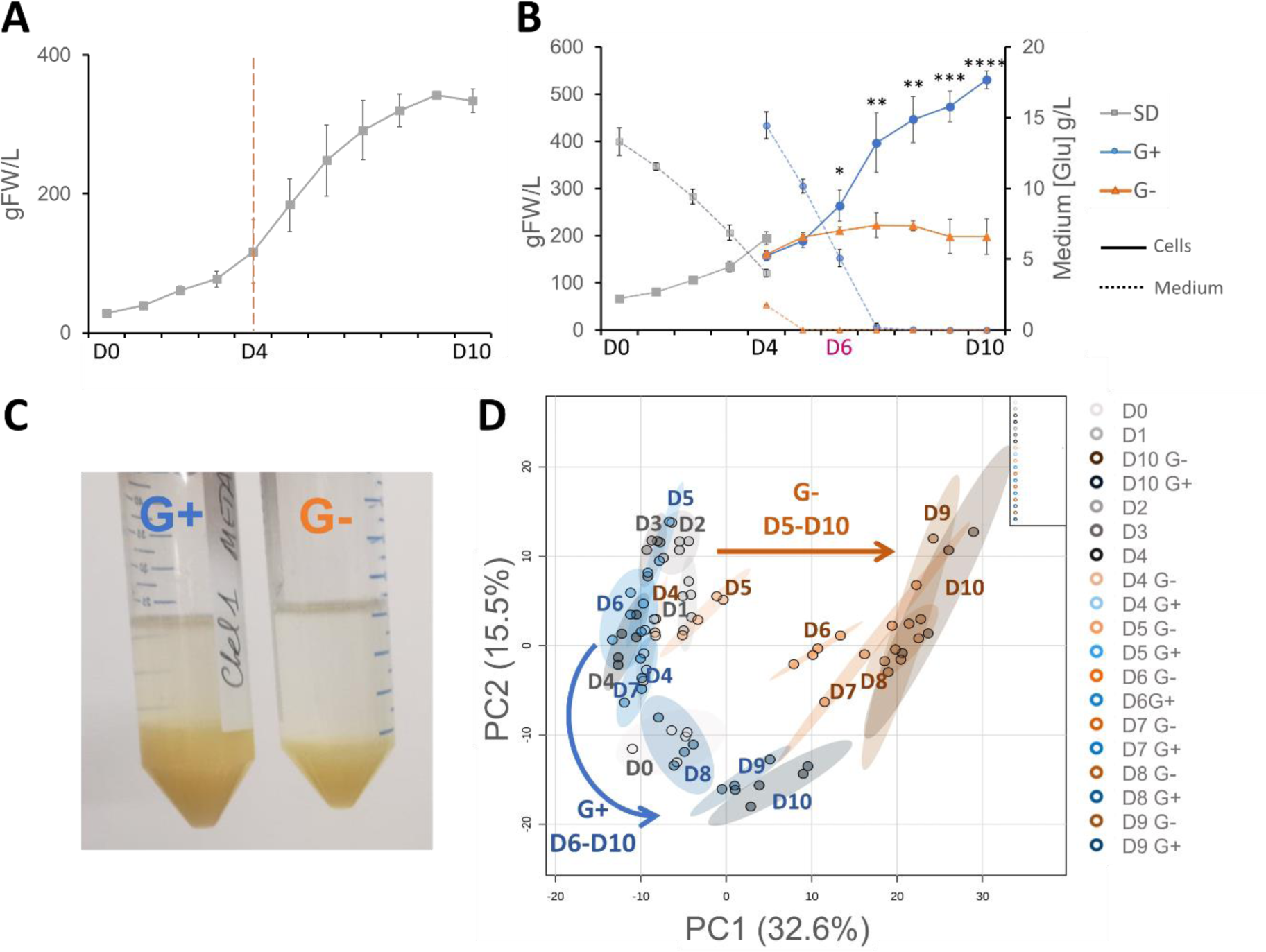
Sugar depletion triggers rapid growth arrest and metabolic drift in grapevine cells. (A) Cell growth curve of Cabernet Sauvignon (CS) cells in standard condition (SD) (n=2), (B) CS cell growth kinetic (solid line) and extracellular glucose consumption (dotted lines) in SD (D0-D4, grey line) and after subculture in glucose rich (G+ blue line) or glucose poor (G-, orange line) from D4 to D10 (n=4). Vertical bars are CI (n ≥ 4). Significant differences between G+ and G- cells are shown (*p ≤ 0.05, **p ≤ 0.01, *** p ≤ 0.001, **** p ≤ 0.0001). (C) G+/G- cells amount 48h after transfert. (D) Principal Component Analysis (PCA) score plots (n=4) of 1719 LCMS-based metabolic signatures under SD, G+ or G- conditions. Maximal variance explained by each PC is given in brackets. Arrows represent the direction of metabolomic evolution in the PCA. SD (D0-D4): grey shades; G+ (D4-D10): blue shades; G- (D4-D10): orange shades.

At D6, G- cells growth was significantly reduced as compared to G+ cells (Figure 1B and C). However, as shown in Figure 1B, the glucose concentration in the medium decreased rapidly in G+ cells. Of note, glucose concentration decreased more rapidly in medium of G+ treated cells (D4 to D10) than in STD (D0 to D4). In the STD condition, extracellular glucose concentration decreased from 13.3 (± 0.97) g/L at D0 to 11.57 (± 0.26) g/L at D1, and from 14.46 (± 0.94) g/L at D4 to 10.16 (± 0.50) g/L at D5. This difference most likely reflect the higher number of cells present at D4 after subculturing the cells in G+ medium (156.7 ± 10.2 gFW/L) as compared to the cell amount at D0 (66.8 ± 7.0 gFW/L), thus increasing extracellular glucose consumption rate.

At D6, the glucose uptake rate was higher in G+ than in G- (5.44g/L/day and 2g/L/day respectively). In G- condition, glucose was no longer detectable at D5 in contrast to G+ conditions whereas glucose was detectable up to D7. At D6, the glucose uptake rate was the highest (5.44 g/L/day) in G+ condition and was null in G- condition. This resulted in different Osmotic Pressure (OP) between conditions, that could be evaluated based on the measured glucose concentration in the medium with a maximum of 110 mOs delta between G+ and G- medium observed at D4 (Figure S2). This difference was however transient and no more detectable from day 7 (Figure S2). As a conclusion, limiting carbon availability to cells leads to a dramatic and rapid reduction of cell growth. Regarding glucose uptake rate, D6 appears as the most contrasted conditions to study grapevine cell response to carbon deficiency.

### Sugar depletion triggers a major metabolic shift in grapevine heterotrophic cells

To characterize the consequences of carbon limitation on grapevine cell metabolism, LCMS analyses were performed on cells harvested each day from D0 to D4, and, after subculture, from D4 to D10 in G+ and G- conditions (n ≥ 4). After filtering the most 1 719 reliable variables (see methods) corresponding to all metabolite signatures quantified considering all samples together. Principal component analysis (PCA) showed that samples were grouped at D4 and started to diverge along the PC1 axis (representing 32.6% of the total variance) at D5 (Figure 1D). Maximum multivariate separation between samples was evident at D6 and D7, whilst G+ cells converged toward G- cells after D8 and this trend consistently aligned with a restriction in glucose availability under G- conditions. The G+ cells were separated along the PC2 axis (15.5% of the total variance) as a function of time. Pearson’s correlation clustering (Figure S3) further confirmed the divergence in separation between samples as influenced by both incubation time and medium composition.

Likewise, pairwise comparisons of the metabolic profiles showed that the number of metabolites (P<0.01) differentially accumulated between G+ and G- conditions increased from 1 and 56 at D4 and D5, respectively, followed by a five-fold increase up to 285 metabolite signatures from D5 to D6, with no further increase at D7 (Table 1). The number of differentially accumulating metabolites reached 433 at D8 before decreasing from 213 to 132 at D9 and D10, respectively (Table 1). To better visualize these changes, metabolomics profiling of G+ and G- cells at D4, D5, D6 and D7 were further used to extract the 596 signals that are the most significantly differentially accumulated in all samples (P<0.01) over the 1703 signals detected from D4 to D7 (Figure S4). Utilizing Pearson’s correlation clustering on the 596 signals resulted in the classification of samples into four distinct groups. These groups exhibited variations in metabolite accumulation profiles from D5 to D7, contingent upon the specific growth conditions, as illustrated in Figure S4.

**Table 1.**
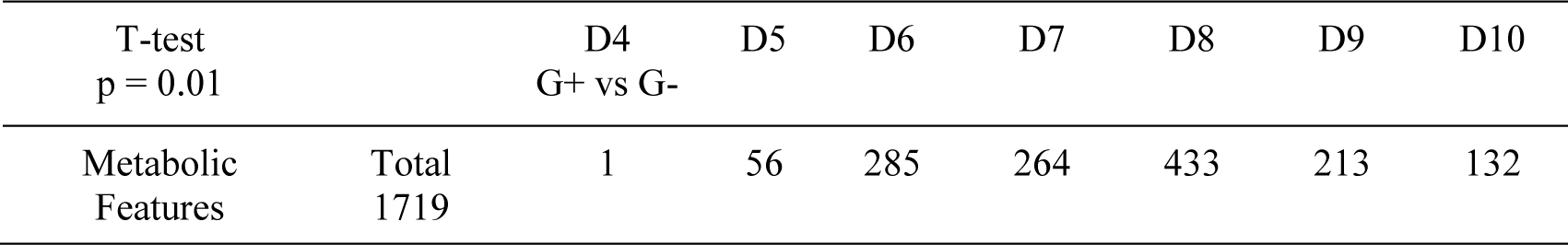
Pairwise statistical analysis of the metabolomic features differing between G+ and G- conditions from D4 to D10.

| T-test<br>p = 0.01 |  | D4<br>G+ vs G- | D5 | D6 | D7 | D8 | D9 | D10 |
| --- | --- | --- | --- | --- | --- | --- | --- | --- |
| Metabolic<br>Features | Total<br>1719 | 1 | 56 | 285 | 264 | 433 | 213 | 132 |

Overall, global metabolome analysis reveals that changes in glucose availability generate major metabolic adjustment in heterotrophic cells as early as 24h after subculture.

### Sugar limitation has a profound impact on the cell primary metabolism and redox state

To obtain a more targeted view of the metabolic consequences generated by carbon shortage, soluble sugars, organic acids, proteins, and total amino acids were quantified from D0 to D10 in all growing conditions (Figures 2A-G, Figure S5). As observed with untargeted metabolic analysis, most of the metabolites showed significant differences in abundance between G+ and G- conditions starting from D6.

**Figure 2.**
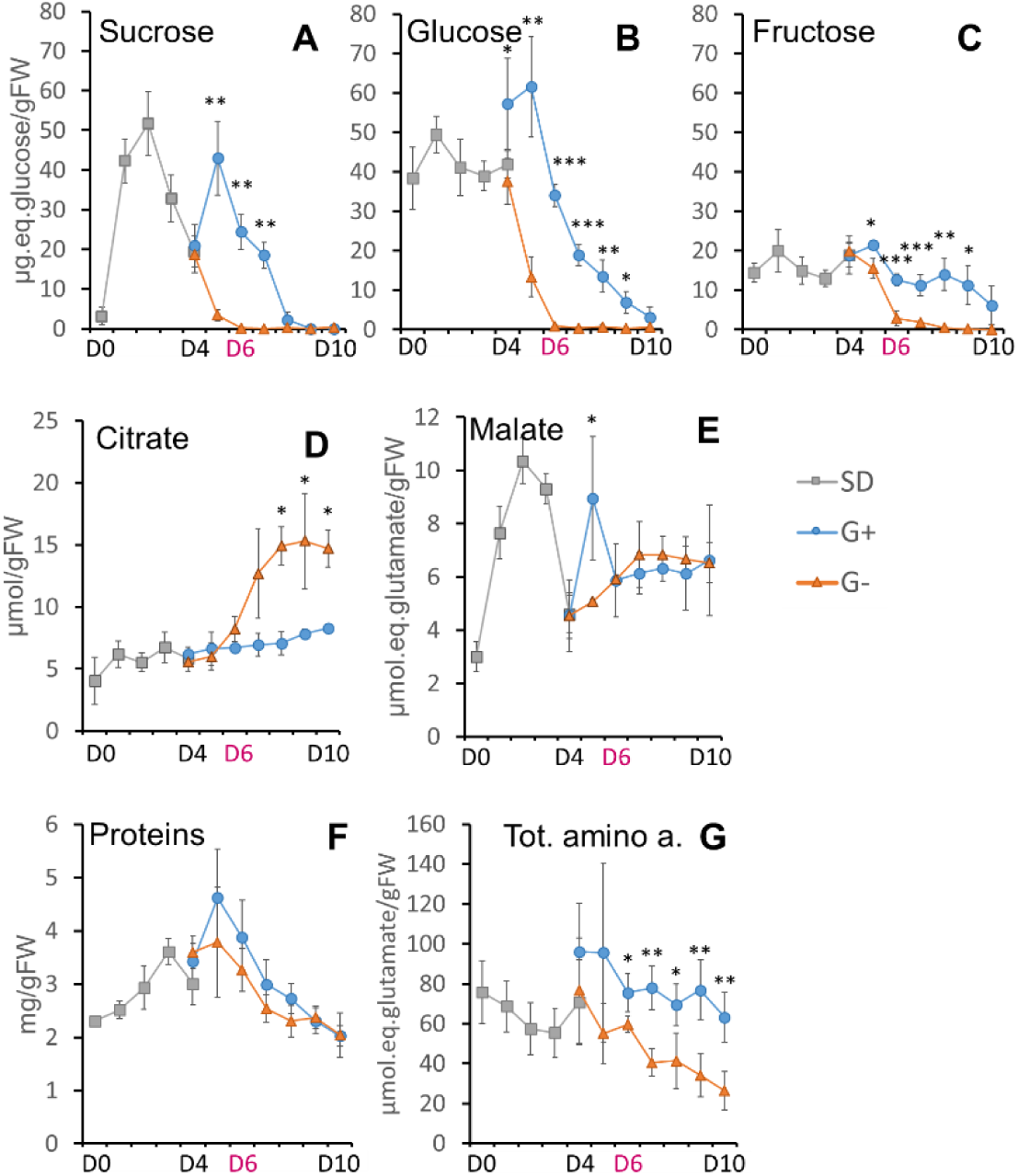
Time-course of targeted primary metabolites accumulation in cells from D0 to D10. Accumulation of soluble sugars (measured in µq.eq.glucose/gFW) (A-C), organic acids (D, E), total protein (F) and amino acids (G). Stars indicate the level of significance of G+ and G- sample comparison (*p ≤ 0.05, **p ≤ 0.01, *** p ≤ 0.001, **** p ≤ 0.0001).

In pre-cultured cells (D0 to D4), the glucose and fructose cell content did not vary significantly (Figure 2B and C), in contrast to sucrose that increases rapidly from D0 (3.13 ± 2.6 µg.eq.glucose/gFW) to peak at D2 (51.76 ± 8.06 µg.eq.glucose/gFW), before decreasing to reach 19.39 ± 1.94 µg.eq.glucose/gFW at D4 (Figure 2A). Although the accumulation of total amino acids gradually decreased from D0 to D4, individual amino acids presented various accumulation profiles (Figure S5). After the transfer of cells to G- condition all soluble sugars rapidly dropped from D4 to reach levels below detection at D6 (Figure 1A-C). In contrast, in cells transferred in G+ conditions, intracellular glucose content rapidly increased to 61.6 ±12.8 µg /gFW at D5 and then decreased following a biphasic kinetic to reach 3.2 ± 2.6 µg/gFW at D10. A different profile is observed for fructose, which presents a weak and transient increase at D5 followed by a progressive decrease to reach a plateau at D6 that is maintained up to D10 at a level similar to SD conditions (Figure 1C). Like glucose, the cell sucrose content increased over 2-fold from D4 to D5 in G+ conditions before decreasing to levels below detection at D8 (Figure 1A, B). To the exception of a transient increase in malate content at D5 in G+ cells (Figure 2E), total protein and malate abundance showed little difference in accumulation profiles between G+ versus G- conditions (Figure 2E, F). In contrast, the abundance in total amino acid contents decreased significantly from D6 on, although amino acids presented five different profiles of accumulation (Figure 2G, supplementary fig 5.). Inversely, citrate started to accumulate since D5n G- condition (Figure 2D).

The NAD(P)H/NAD(P) ratio (reduced/ oxidized forms) a good indicator of the redox state of cells (W. Xiao et al., 2018). Both NAD(H) and NADP(H) were measured, and their reduced/oxidized ratio calculated (Figure 3). The NADPH/NADP and NADH/NAD ratio did not show significant variations during the preculture (D0 to D4). However, after subculturing, the NADPH/NADP ratio was highly variable in both conditions from D5 to D7. At later times, it increased in G+ cells to reach 1.18 ± 0.58 at D10, twice more than the value of 0.58 ± 0.22 calculated in G- conditions. The NADH/NAD ratio showed a significant increase, from D6 (0.07 ± 0.04) to D10 (0.18 ± 0.02) in G+ condition and an important decrease in G- condition, to reach a final value below 0.1 (Figure 3). Hence glucose limitation impacts the redox balance in grapevine cells, with a progressive decrease of the oxidized relative to reduced form of both NAD and NADP.

**Figure 3.**
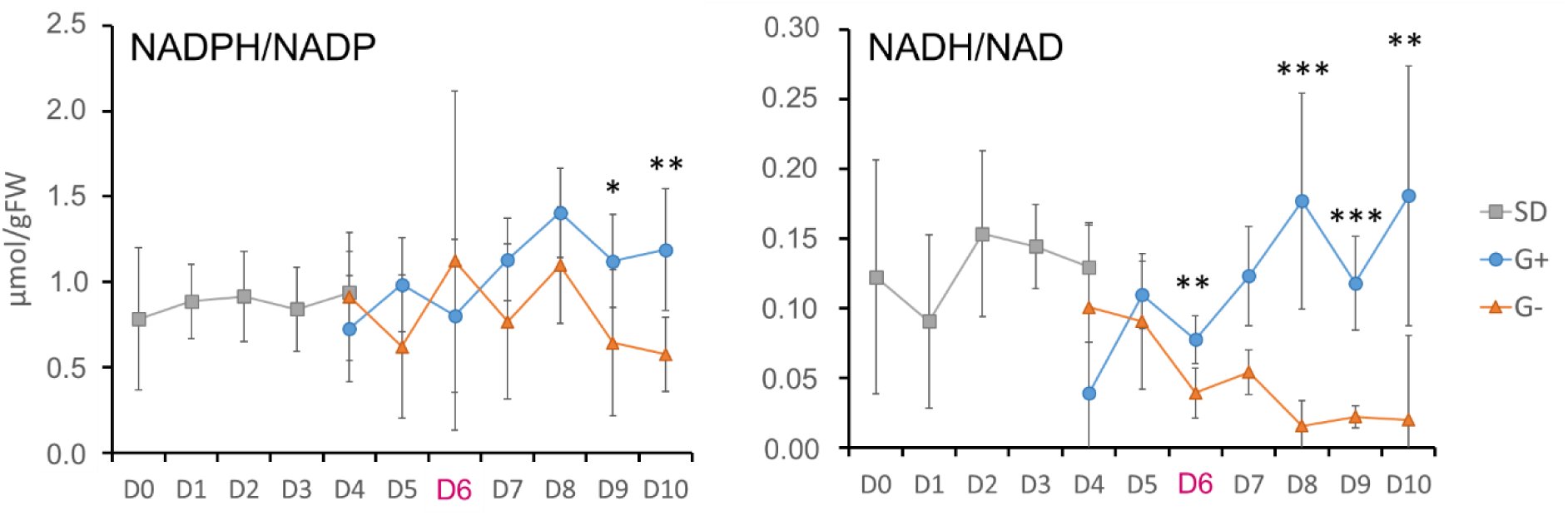
Time-course of NAD(P) oxidized and reduced form proportion through calculation of NAD(P)H/NAD(P) ratio in SD (grey), G+ (blue), G- (orange) conditions. Vertical bars represent CI (n=8). Stars indicate the level of significance of G+ and G- sample comparison (*p ≤ 0.05, **p ≤ 0.01, *** p ≤ 0.001).

Overall, G- condition strongly affects soluble sugars accumulation in cells, but also impacts the abundance of many other metabolites, including free amino acids, organic acids, or total proteins, and affects the NAD(P)H/NAD(P) balances consistent with a strong impact of carbon deprivation on grapevine cell central metabolism, and on the redox state.

### Flux Balance Analysis reveals that carbon limitation alleviates cell metabolic fluxes and impacts the 1C metabolism in grapevine cells

The experimental data of the targeted metabolites and cell components were integrated to constrain a flux model analysis and evaluate the impact of carbon limitation on the metabolic fluxes of cells. To perform the flux analysis (Table S1-S4), the metabolic model was constrained to calculate fluxes as a snapshot each day in all conditions (*i.e*. in the standard culture (SD) and in G+ and G- conditions). Because of remaining glucose in the medium at D4 in G+ condition and of limiting glucose after D7 in both conditions, we focused on D6 to evaluate the consequences of change in carbon availability on the metabolic fluxes and to integrate results with those of molecular analyses. We were mainly interested in the changes of fluxes as a way to evaluate metabolism reprogramming under carbon starvation. The comparison of fluxes (Table S5) shows that the main changes in external fluxes (used as constraints in the metabolic model) in G- is an increase of fluxes toward amino acids synthesized from oxaloacetate (OAA) and a decrease of those of cell wall synthesis and hexoses storage.

Most of the internal fluxes calculated in G- were lower than the ones calculated in G+ at D6 (Figure S6) including those of glycolysis, oxidative phosphate pathway, and cell wall biosynthesis, and to a lower extend those of respiration, nitrate assimilation, and of the TCA cycle. Interestingly, some fluxes were increased under carbon limitation. They include fluxes involved in the mobilization of stored compounds, such as cell wall, protein, as well as the accumulation of lipids, and of some amino acids. This led to an increase of internal fluxes mainly through fructokinase, malate dehydrogenase, malic enzyme, aminotransferases (aspartate and alanine) required to generate the precursors to support the synthesis and accumulation of amino acids and organic acids and the redox fluxes towards ascorbate and glutathione accumulation. Consistent with the requirement of organic acid and amino acid storage, a higher flux partition is observed towards the anaplerotic pathway (*Vpepc/Vpepc+Vpk*) ratio higher in G- (0.371) than in G+ (0.293) at D5). These flux results show a clear metabolic stress for cells under carbon limitation.

We focused on the folate and methionine cycles (the 1C metabolism), as they link primary metabolism to methylation (Lindermayr et al., 2020). For the THF and SAM cycles, we used the calculated fluxes to estimate the flux activity for an intermediate metabolite as the sum of fluxes producing (or consuming) this metabolite (Figure 4, Table S6). Most of the fluxes were rapidly reduced down to zero between D4 and D6 under G-, except fluxes through dihydrofolate (DHF) and in less extent through tetrahydrofolate (THF) and 10-formyltetrahydrofolate (fTHF), showing 0.20×10^-3^ mmol/g/DW/day in G- condition as compared to 0.85 10^-3^ mmol.g.DW^-1^.day-1 in G+ conditions (Figure 4). By contrast, the SAM fluxes estimated at 0.075 mmol/g/DW/day in G+ condition at D6 were reduced to 0.012 10^-3^ mmol/g/DW/day in G- condition (Figure 4, Table S6). This suggests that folate synthesis is not completely inhibited under carbon limitation, but that parts of this cycle are prioritized as compared to others. According to model prediction, folate is *a priori* mainly needed for other DHF-dependent pathways such as nucleotides synthesis (Table S2), rather than for DNA methylation, because SAM-cycle fluxes are decreased under carbon limitation. Overall, these results show that sugar limitation impacts 1C metabolic pathways, including those implicated in the synthesis of SAM.

**Figure 4.**
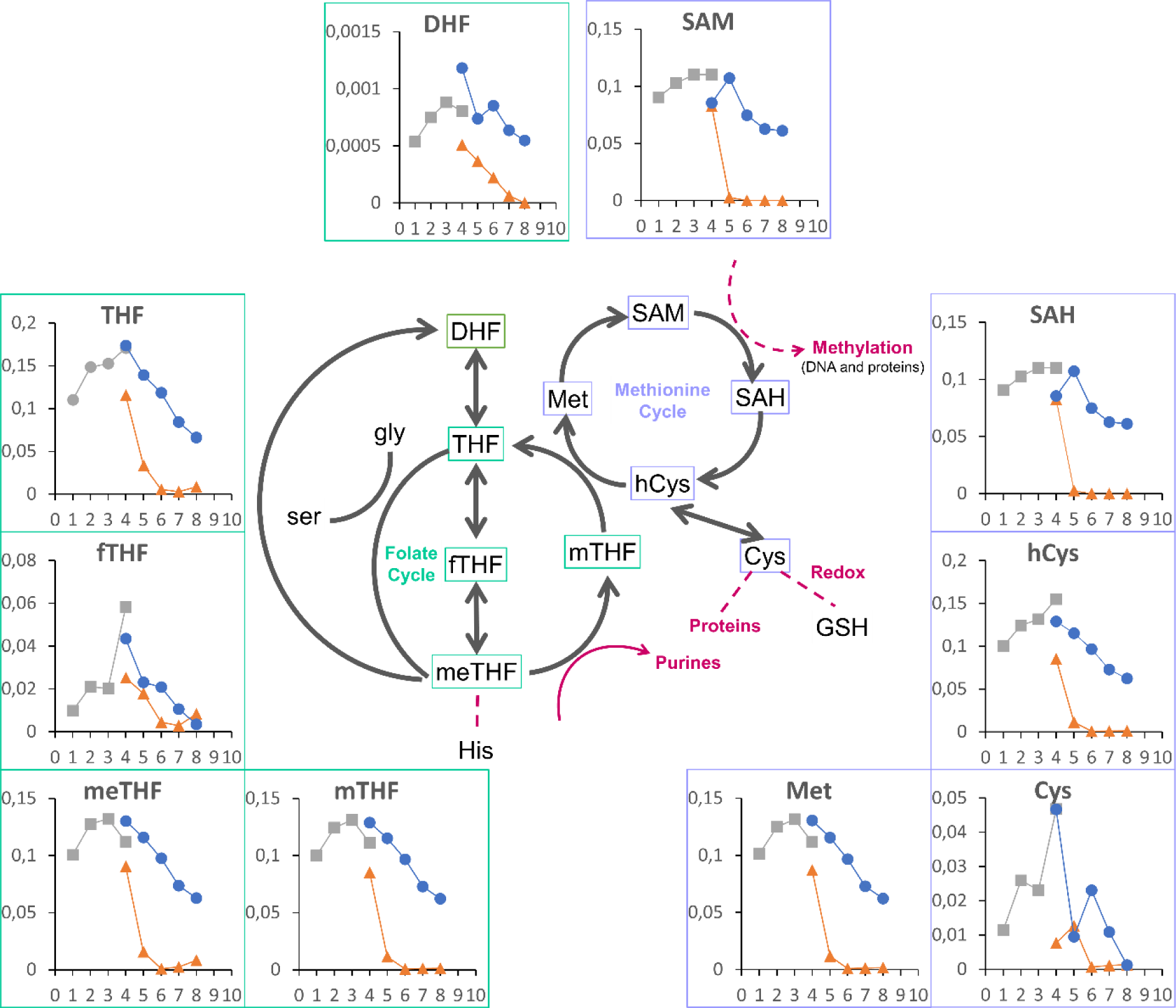
Graphic representation of flux kinetical analysis of TetraHydroFolate (THF) and S-Adenosyl-Methionin (SAM) cycle. The x-axis represents the Days, the y-axis represents flux values measured in mmol.gDW-1.Day-1. Grey lines from D1 to D4: SD condition; from D4 to D8 blue line: G+ cells; from D4 to D8 blue line: G- condition.

### Genes of several metabolic pathways display differential expression under sugar depletion

To characterize the transcriptomic response of CS cells to glucose depletion, RNAseq analyses were performed at D6 in both G+ and G- conditions. After trimming, quality assessment and removal of low-quality reads, total cleaned reads, approximately ∼24 million reads for each sample, were mapped to the grapevine reference genome (12X.V2; Canaguier et al., 2017), of which on average 90% were uniquely mapped (Table S7). Between 69.7% and 71% of aligned reads overlap known genes. PCA analysis performed using all expressed genes reveals both strong differential behavior between G+ and G- conditions, and low variability between biological replicates (Figure S7). This observation is also supported by the analysis of the 200 genes showing the highest variation in expression levels between conditions (Figure 5). Replicates for each condition cluster together in correlative analysis, but are clearly separated between G+ and G- conditions.

**Figure 5.**
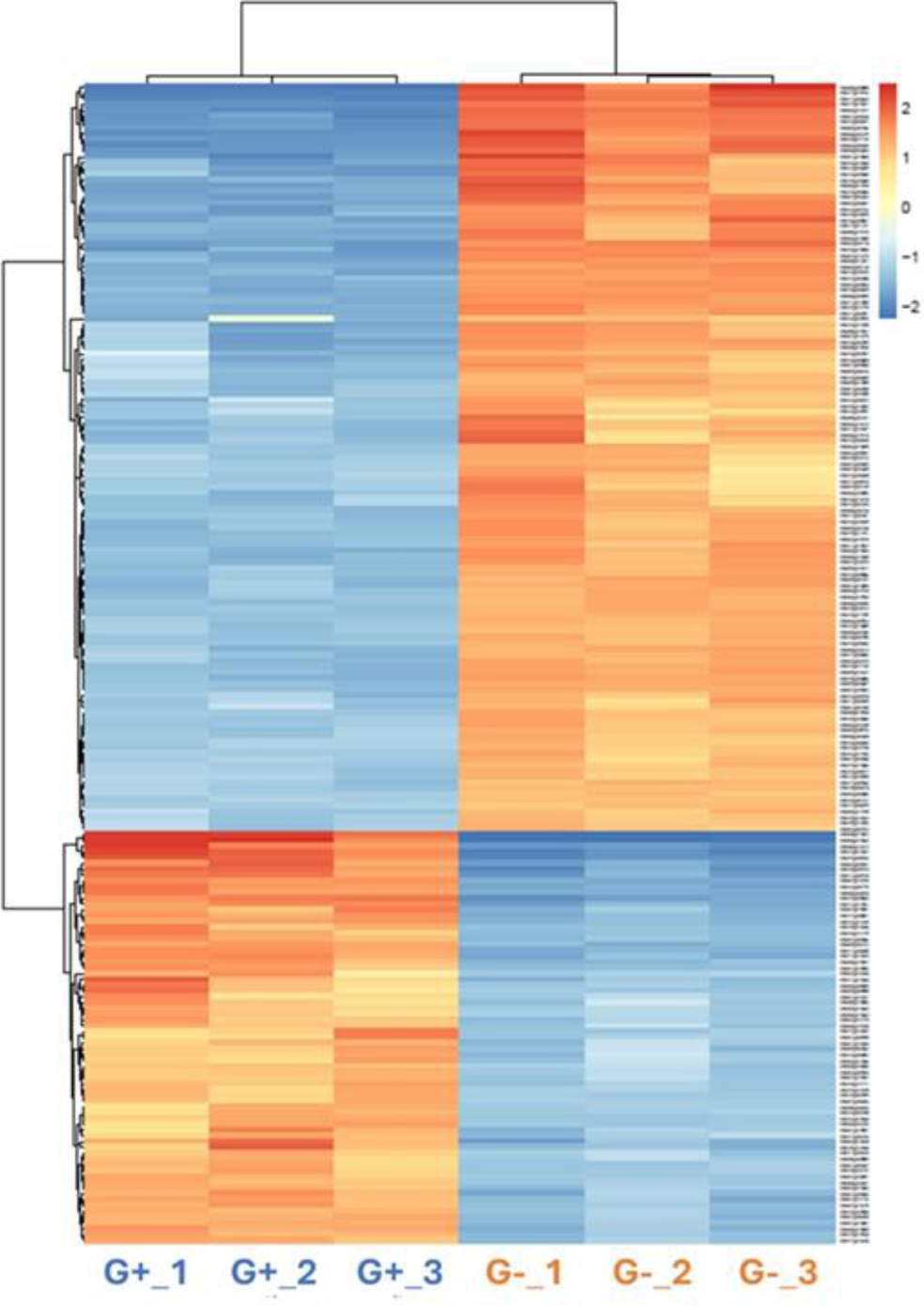
Heatmap representation of normalized expression of the 200 genes showing the highest variation within G+ and G- samples analyzed at D6 (pval <0.05) Colors represent the variability of gene expression among samples, with a normalization per row. Dendrogram were calculated using pearson’s correlation. G+: cells cultivated in glucose rich medium, G-: cells cultivated in glucose poor medium. Mapman visualization of identified DEGs in grapevine in G- cells at D6.

After removal of the low count genes, a total of 5 607 genes were considered differentially expressed (Log2FoldChange >1.0; padj 0.05), among which 2 802 were upregulated and 2 805 downregulated in G- versus G+ conditions. A gene ontology (GO) overrepresentation analysis was performed to identify the biological processes affected by carbon deprivation. For upregulated genes, “photosynthesis processes” including light harvesting, chlorophyll biosynthesis and electron transport in photosystems were overrepresented, together with genes involved in stress responses (hormones, redox process, defense response), and in metabolite biosynthesis and transport (trehalose, malate, oligopeptide; Figure S8). By contrast, downregulated genes were enriched in genes involved in translation (ribosome biogenesis, translational elongation), cell division (DNA replication, microtubule-based movements, chromosome segregation), amino acids biosynthesis (serine, threonine, glycine, tyrosine, phenylalanine), and processes related to carbon metabolism (gluconeogenesis, glycolysis, sugars, fatty acids; Figure S8).

Using a MapMan representation with both targeted metabolites and RNAseq data (Figure 6A and B), 89 and 947 out of 5 607 DEGs were successfully assigned to cell cycle and various metabolic pathways respectively. A majority of DEGs related to the regulation of cell cycle (82/89) were downregulated at D6 (Figure 6A), suggesting that the cell growth arrest observed in G- conditions correlates with a stop of cell divisions. When considering DEGs involved in metabolism (Figure 6B), those associated with cell wall construction (pectin esterases, expansins and xyloglucane endotransglycosylases (XETs)), in amino acids and nucleotide synthesis, glycolysis, TCA cycle, energy metabolism (Mitochondrial Electron Transport), sucrose and starch synthesis, in glycolysis and other downstream reactions were downregulated in G- conditions (Figure 6B, Figure S9). Inversely, DEGs associated with the mobilization of carbon resources such as amino acids, lipids, and sucrose, starch and protein breakdown together with the glyoxylate cycle were upregulated (Figure 6B, Figure S9). Even though cells are non-photosynthetic and heterotrophic, genes involved in the light reactions of photosynthesis were strongly upregulated in G- cells compared to G+ ones (Figure 6B), as well as those of in photosystems maintenance and electron transport chain reactions (Figure 6B, Figure S10).

**Figure 6.**
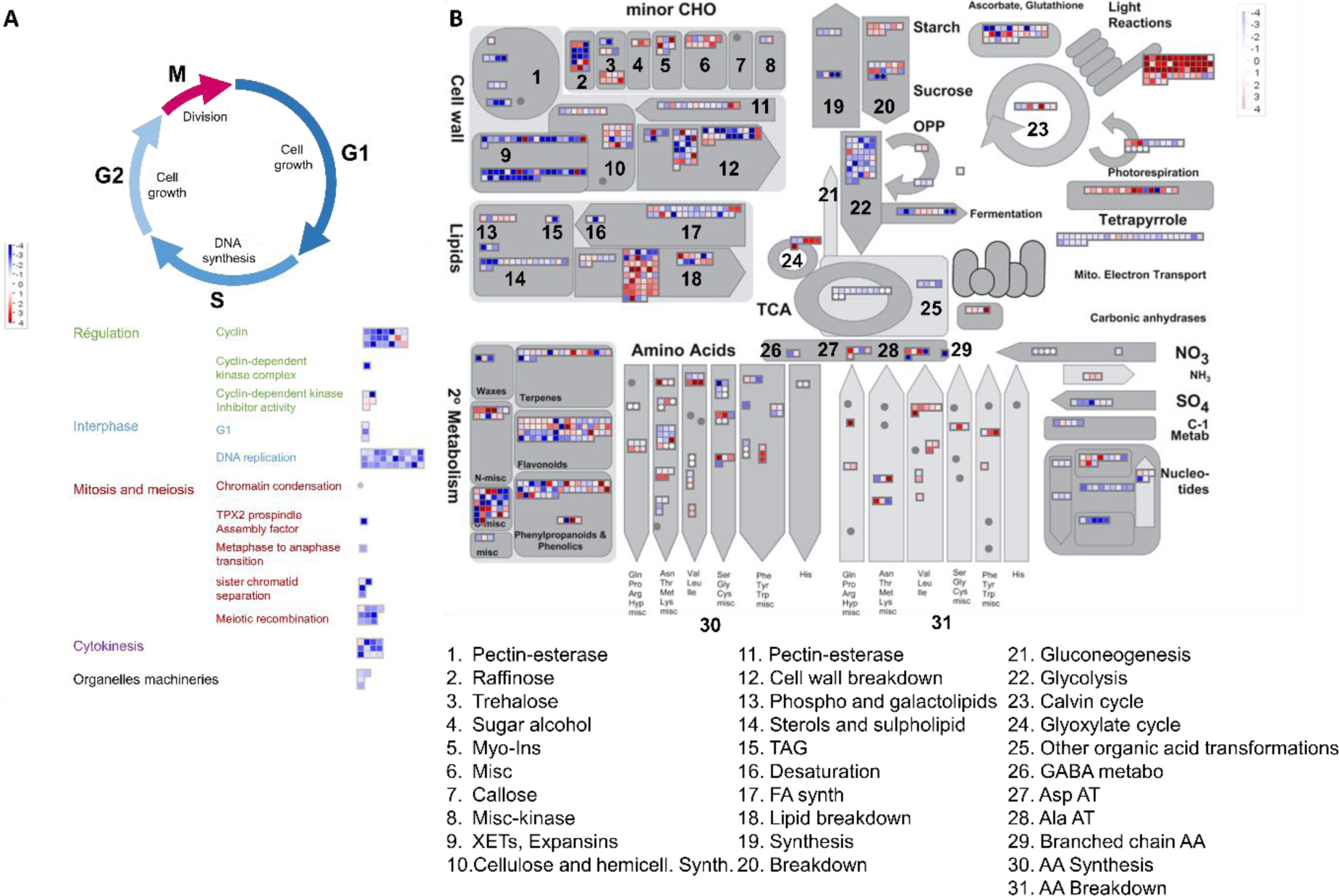
Mapman visualization of identified DEGs in grapevine in G- cells at D6. (A) Cell cycle-related genes. Each square represents a gene expressed at a higher level in G- cells (red) or in G+ cells (blue). (B) Metabolism overview Mapman representation obtained from the DEGs and targeted metabolites analysis in G- cells at D6. Each square represents a gene, each circle represents a metabolite. In red: genes more expressed in G- cells; in blue: genes more expressed in G+ condition.

Genes involved in 1C metabolism were downregulated at D6 (Figure 6), including those related to the synthesis of methylenetetrahydrofolate (methylene-THF), a precursor of the tetrahydrofolate cycle (THF cycle) (Figure S10). Overall, results show that sugar deprivation strongly impacts the expression of genes of the metabolic pathways dedicated to carbon mobilization, nutrient mobilization, and heterotrophic carbon formation and are consistent with previous data obtained on Arabidopsis cells under sugar starvation (Contento et al., 2004).

### Genes involved in epigenetic processes are differentially regulated under carbon limitation

To investigate possible epigenetic regulations during carbon deprivation we analyzed the expression of genes involved in epigenetic processes in G+ and G- cells. We first assessed the expression of DNA-methylation modifiers by investigating RNAseq data with ID of grapevine DNA methyltransferases and demethylases (Junhua Kong, PhD Thesis). Three DNA-methyltransferases (DNMTs), VvCMT1 (*Vitvi08g01767*), VvCMT3 (*Vitvi06g00102*), VvCMT4 (*Vitvi16g00174*), one DNA-demethylase (DML) VvDML3 (*Vitvi06g01402*), and one helicase VvDDM1 (*Vitvi04g01275*), were differentially expressed in G- condition as compared to control (Table 2). The four DNMTs were all down-regulated with a Log2FoldChange (L2FC) ranging between -1.06 and -1.45 (padj<0.01). Similarly, VvDML3 was repressed with a L2FC of -2.60 (padj<0.01).

**Table 2.:**
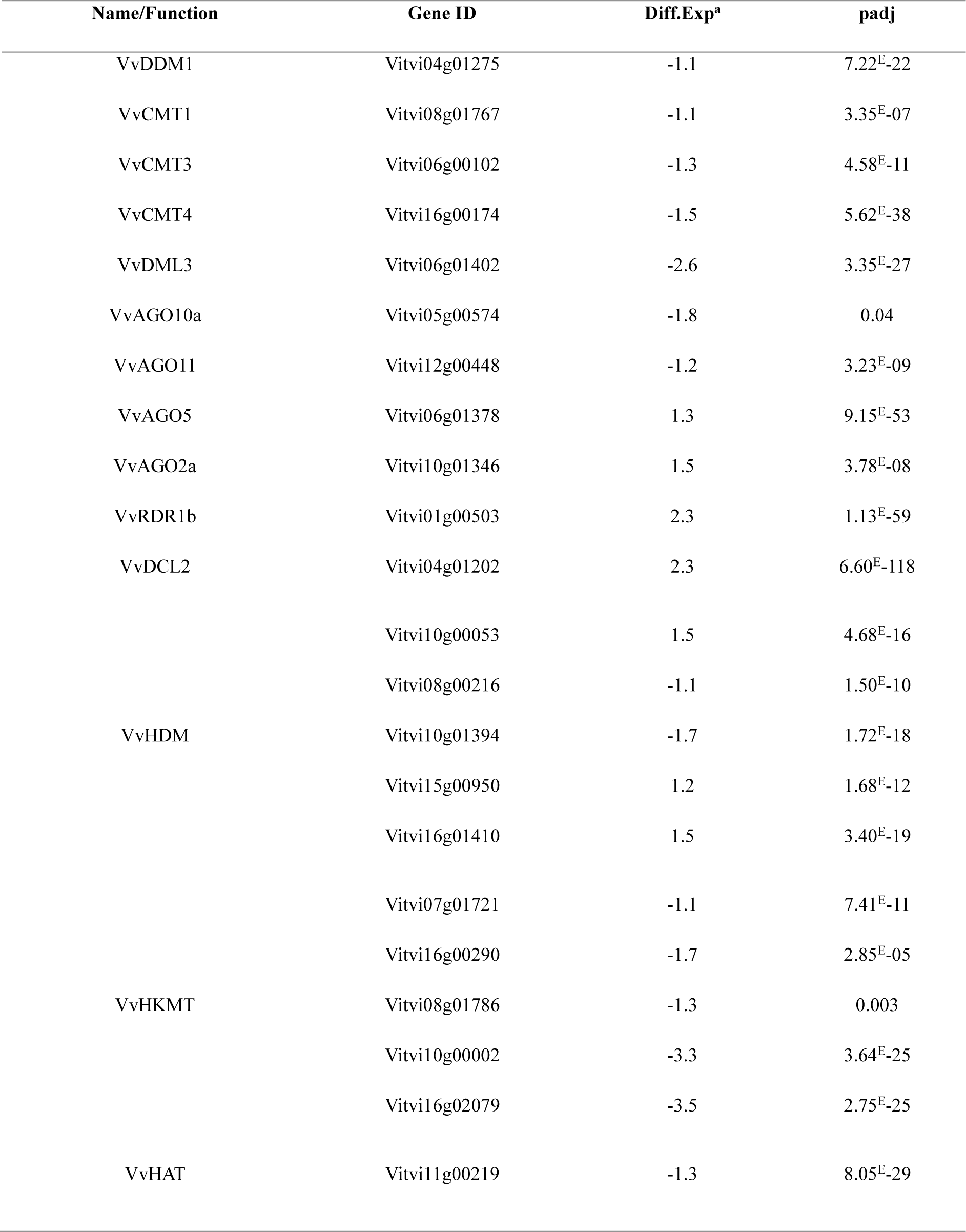
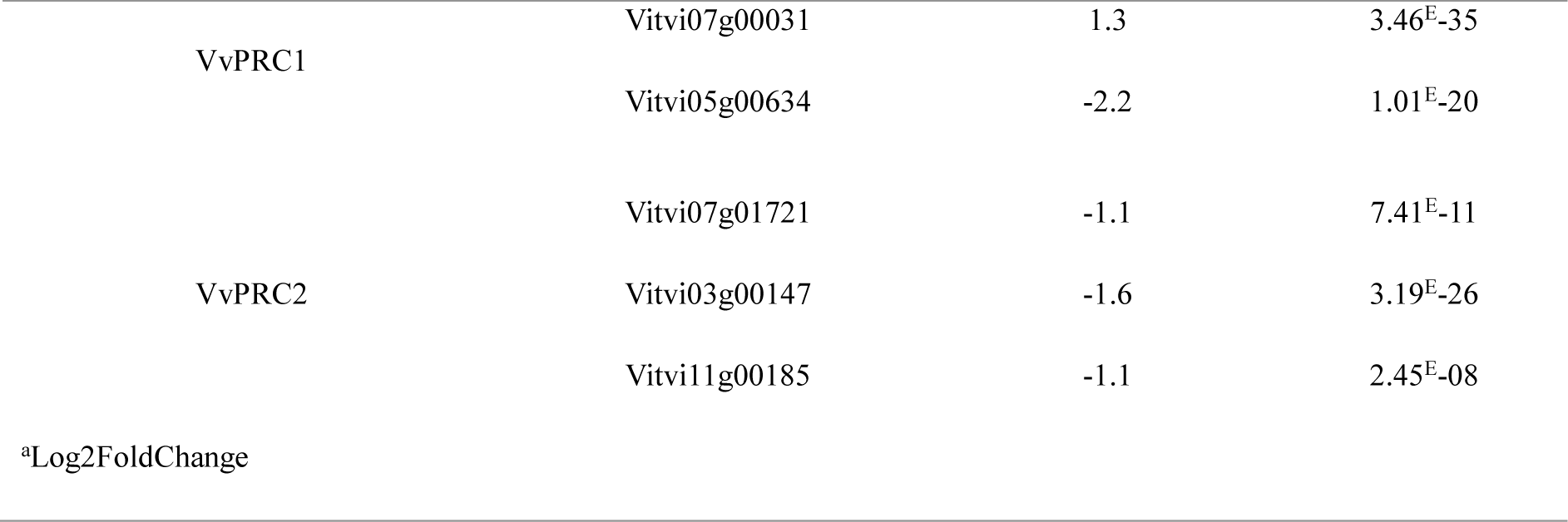
Grapevine epigenetic modifiers differential expression in G- cells. DDM1: DECREASED DNA METHYLATION 1; CMT: CHROMOMETHYLASE; DML: DEMETER-LIKE; AGO: ARGONAUTE; RDR : RNA-DEPENDENT RNA POLYMERASE; DCL: DICER-LIKE; HDM: Histone Demethylase; HKMT: Histone-Lysine-Methyl-Transferase; HAT: Histone Acetyl-Transferase; PRC: Polycomb Repressive Complex.

We also analyzed genes involved in siRNA synthesis, as they may play a critical role in DNA methylation control. The expression of grapevine Dicer-Like (DCL), Argonaute (AGO), and RNA-dependent RNA Polymerase (RDR) using (Zhao et al., 2015) work as a reference (Table 2). Four ARGONAUTE (VvAGO10a, VvAGO11, VvAGO5, VvAGO2a), one RNA-Dependent-RNA polymerase (VvRDR1b) and one DICER-like (VvDCL2) protein were found differentially expressed under G- condition. VvAGO10a and VvAGO11 were both down-regulated showing L2FC values of - 1.8 and -1.2 respectively (padj<0.05), while the four other genes were up-regulated with a L2FC of 1.3 for VvAGO5 and 1.5 for VvAGO2a (padj<0.05) and a L2FC of 2.3 for both VvRDR1b and VvDCL2 (padj<0.05) (Table 2).

The RNAseq data were further investigated using the ID list of recently identified histone modifiers in grapevine (Li Wang et al., 2020) (Table 2), and POLYCOMB REPRESSING COMPLEX2 (PRC2) related putative proteins that we have identified (Table 2, Figure S11, Figure S12). Five histone demethylases (HDMs, *Vitvi10g00053*, *Vitvi08g00216*, *Vitvi10g01394*, *Vitvi15g00950*, *Vitvi16g01410*), and histone lysine methyltransferases (HKMTs, VvCLF/*Vitvi07g01721*, *Vitvi16g00290*, *Vitvi08g01786*, *Vitvi10g00002*, *Vitvi16g02079*) were differentially expressed under G- condition (Table 2). In addition, one histone acetyltransferase (HAT, *Vitvi11g00219*), two proteins related to the POLYCOMB REPRESSING COMPLEX1 (PRC1) VvDRIP1 (*Vitvi07g00031*) and VvDRIP3 (*Vitvi05g00634*) and to the PRC2, VvMSI2 (*Vitvi03g00147*), VvMSI5 (*Vitvi11g00185*) were also differentially expressed under carbon limitation (Table 2). The five HKMT were all downregulated under sugar limitation with a L2FC distributed between -1.08 and -3.54 (padj <0.05), while HDMs showed mixed expression patterns with two genes being downregulated (logf2FC of -1.09 for *Vitvi08g00216* and -1.66 for *Vitvi10g01394*), and three upregulated genes with a L2FC comprised between 1.18 and 1.53 (padj<0.05). HAT appeared downregulated in carbon limitation condition with a L2FC of -1.31 (padj<0.05), as well as the putative PRC2 proteins VvMSI2 and VvMSI5 with a L2FC of -1.58 and -1.11 respectively (Table 2). The two putative PRC1 proteins identified showed opposite behavior, with VvDRP1 being upregulated (L2FC=1.31, padj<0.05) and VvDRIP3 downregulated (L2FC=-2.20, padj<0.05).

Taken together, these results show that carbon limitation impacts the expression of genes encoding epigenome modifiers, including DNMTs, demethylases and DDM1, histone modifiers, and putative polycomb group proteins, as well as genes related to the Post Transcriptional Gene Silencing (PTGS) and RdDM pathways.

### DNA-methylation landscape changes associated with carbon limitation

To characterize the consequences of carbon limitation on the DNA-methylation landscapes, Whole Genome Bisulfite Sequencing (WGBS) was performed at D6 in both G+ and G- conditions. A minimum of ∼93 million reads were generated for each replicate (Table 3). The sequencing depth after filtering low-quality reads varied between ∼25X and ∼74X depending on samples (Table 3). Hierarchical clustering reveals that replicate cluster together according to their glucose supplementation state (Figure S13).

**Table 3.** Whole Genome Bisulfite Sequencing mapping results and average depth coverage.

| Sample | Sequence pairs analyzed | Nbr. Of paired-end alignments with a unique best hit | Mapping efficiency (%) | Non-uniquely mapped sequences | Average depth coverage |
| --- | --- | --- | --- | --- | --- |
| G+_1 | 130,472,479 | 82,944,788 | 63.6 | 13,213,594 | 33.97668 |
| G+_2 | 282,099,730 | 181,683,529 | 64.4 | 29,427,292 | 74.39895 |
| G+_3 | 139,655,324 | 91,667,849 | 65.6 | 14,282,757 | 37.54222 |
| G-_1 | 96,857,550 | 59,516,335 | 61.4 | 9,877,183 | 24.38944 |
| G-_2 | 93,098,553 | 61,901,086 | 66.5 | 10,136,166 | 25.35674 |
| G-_3 | 99,426,877 | 61,857,519 | 62.2 | 10,945,369 | 25.33656 |

The G- cells showed methylation levels of ∼50.0%, ∼34.6% and ∼4.0% in the CG, CHG and CHH sequence contexts respectively, which are higher than those of G+ cells, respectively ∼47.6%, ∼31.7% and ∼3.2% (Figure 7A, Table 4, Table 5). Hence global methylation levels were significantly higher in cells under carbon restriction than in control cells (Figure 7A, Table 4, Table 5).

**Figure 7.**
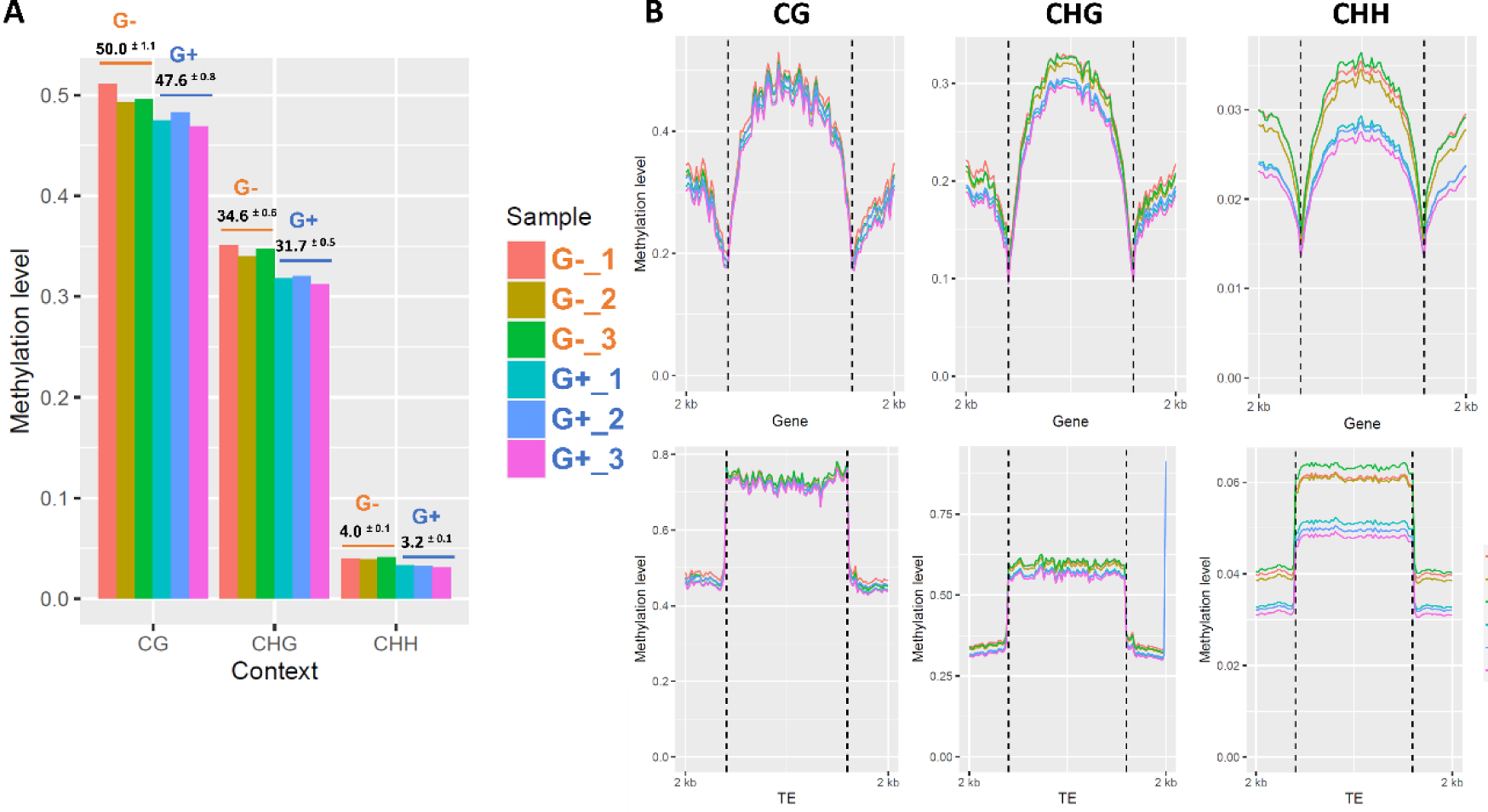
Global methylation levels analysis in G- and G+ condition. (A) Bart chart representation of individual sample methylation levels values (y-axis, 1=100%) in the three-cytosine context (x-axis). (B) Representation of methylated cytosine within and 2kb up- and downstream genes coding sequence.

**Table 4.** Bisulfite sequencing data report.

| Sample | Total of C's analyzed | % of methylated C | % of conversion | Methylated cytosines in |  |  |  | % Methylated cytosines in |  |  |
| --- | --- | --- | --- | --- | --- | --- | --- | --- | --- | --- |
| D6 |  |  |  | CpG | CHG | CHH | Unknown | CpG | CHG | CHH |
| G+_1 | 259,133,083 | 0.567123265 | 99.43287674 | 284,782 | 263,603 | 920,187 | 1,032 | 47.6 | 32.0 | 3.3 |
| G+_2 | 546,752,409 | 0.564586081 | 99.43541392 | 608,384 | 558,261 | 1,918,117 | 2,126 | 48.3 | 32.0 | 3.2 |
| G+_3 | 314,771,657 | 0.601212008 | 99.39878799 | 355,313 | 325,031 | 1,210,998 | 1,103 | 47.1 | 31.4 | 3.1 |
| G-_1 | 159,367,452 | 0.626294759 | 99.37370524 | 199,134 | 180,869 | 617,520 | 587 | 51.2 | 35.2 | 4.0 |
| G-_2 | 168,998,804 | 0.497389911 | 99.50261009 | 179,576 | 159,608 | 500,726 | 673 | 49.5 | 34.1 | 3.9 |
| G-_3 | 162,348,402 | 0.614100901 | 99.38589910 | 208,515 | 190,078 | 597,758 | 632 | 49.8 | 34.8 | 4.1 |

**Table 5.** Methylation average calculted in the three contexts (%), error values correspond to CI (n=3).

| % Methylation | CG | CHG | CHH |
| --- | --- | --- | --- |
| D6_G+ | 47.6 ± 0.8 | 31.7 ± 0.5 | 3.2 ± 0.1 |
| D6_G- | 50.0 ± 1.1 | 34.6 ± 0.6 | 4.0 ± 0.1 |

Analysis of DNA methylation profiles along the entire transcriptional units (gene and 2 kb up or downstream regions) and Transposable Elements (TE) revealed contrasted context-dependent methylation variations (Figure 7B). Cytosine methylation in the CG context remained essentially unchanged between G- and G+ cells. In contrast CHH methylation was enriched up/downstream and within genes in G- cells. Cytosine methylation in the CHG context showed an intermediary behavior with a slight enrichment in G- to compared to G+ cells (Figure 7B). Differentially Methylated Cytosines (DMCs) and Differentially Methylated Regions (DMRs) were determined between G- and G+ cells (Figure 8). A total of 12 524 differentially methylated cytosines were identified between G- and G+conditions with 4 102, 4 307 and 4 115 differentially methylated cytosines (DMCs) in CG, CHG and CHH contexts respectively (Figure 8). DMCs identified in CG context were similarly distributed between hypo and hypermethylated status, while larger proportions of hyper-methylated cytosines were identified in CHG contexts and over 94% in the CHH context (Figure 8).

**Figure 8.**
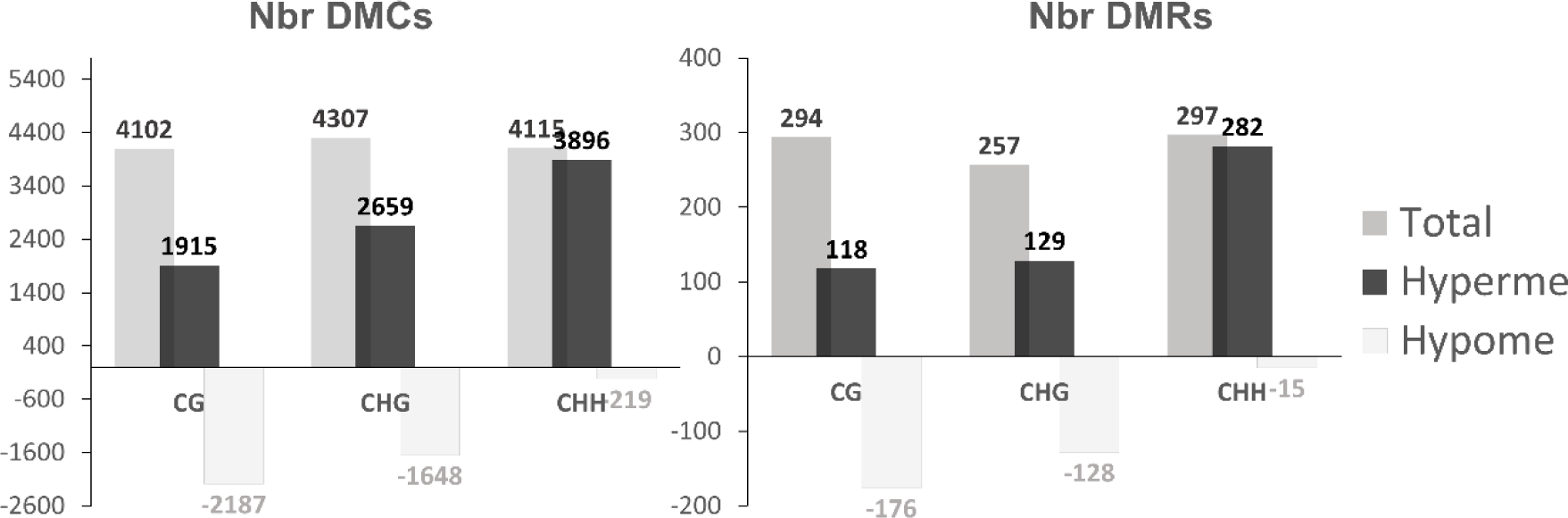
Methylation changes induced by carbon limitation. Identification of Differentially Methylated Cytosines (DMCs) and Differentially Methylated Regions (DMRs) in each context in G- condition compared to G+. Grey bar chart represents the total of DMC/DMRs identified. Direction of methylation changes are represented in black when methylation is gained in G- cells compared to G+ cells (Hyperme) and in light grey when methylation is lost in G- cells compared to G+ cells (Hypome). Number of DMCs and DMRs are indicated on the top/bottom of each barchart.

A similar trend was found when DMRs were analyzed. A total of 848 DMRs were identified in G- cells compared to G+ (Table S8), with 294 CG, 257 CHG and 297 CHH DMRs (Figure 8). Hypo-DMRs are more abundant than hyper-DMRs in the CG context (176 hypo-DMRs over 294 DMRs= 59%), but the converse is found in the CHH context (282 over 297; 95%; Figure 8). By contrast, DMRs in the mCHG context were equally distributed between hypo- and hyper-methylated states (Figure 8). Among the DMRs, 58% (482 over 848) were localized in extragenic Transposable Elements (TEs) and repeats and 22% (186 over 848) in promoter (promDMRs; Figure 9, Table S9). The last 20% (165 over 848) are intragenic-DMRs, distributed between exonic (7%, 59 over 848) and intronic (13%, 106 over 848) regions. The majority of intronic DMRs (75 over 106) and promDMRs (168 over 186) are overlapping with TE inserted upstream or within genes (Figure 9), suggesting that changes in DNA-methylation in response to carbon limitation preferentially targets TE. Non-TE-promDMRs and intragenic-DMRs were mostly found in the and CHG contexts (Figure 10), in contrast to the overrepresentation of CHH DMRs in extragenic regions and TE-promDMRs (Figure 9). The proportions of hyper- and hypo-DMRs were dependent on their localization in the genome. Intragenic-DMRs and non-TE-promDMRs were mostly hypo-methylated in CG and CHG context, while the majority of TE-promDMRs and TE-DMRs were hyper-methylated in the CHH context (Figure 9).

**Figure 9.**
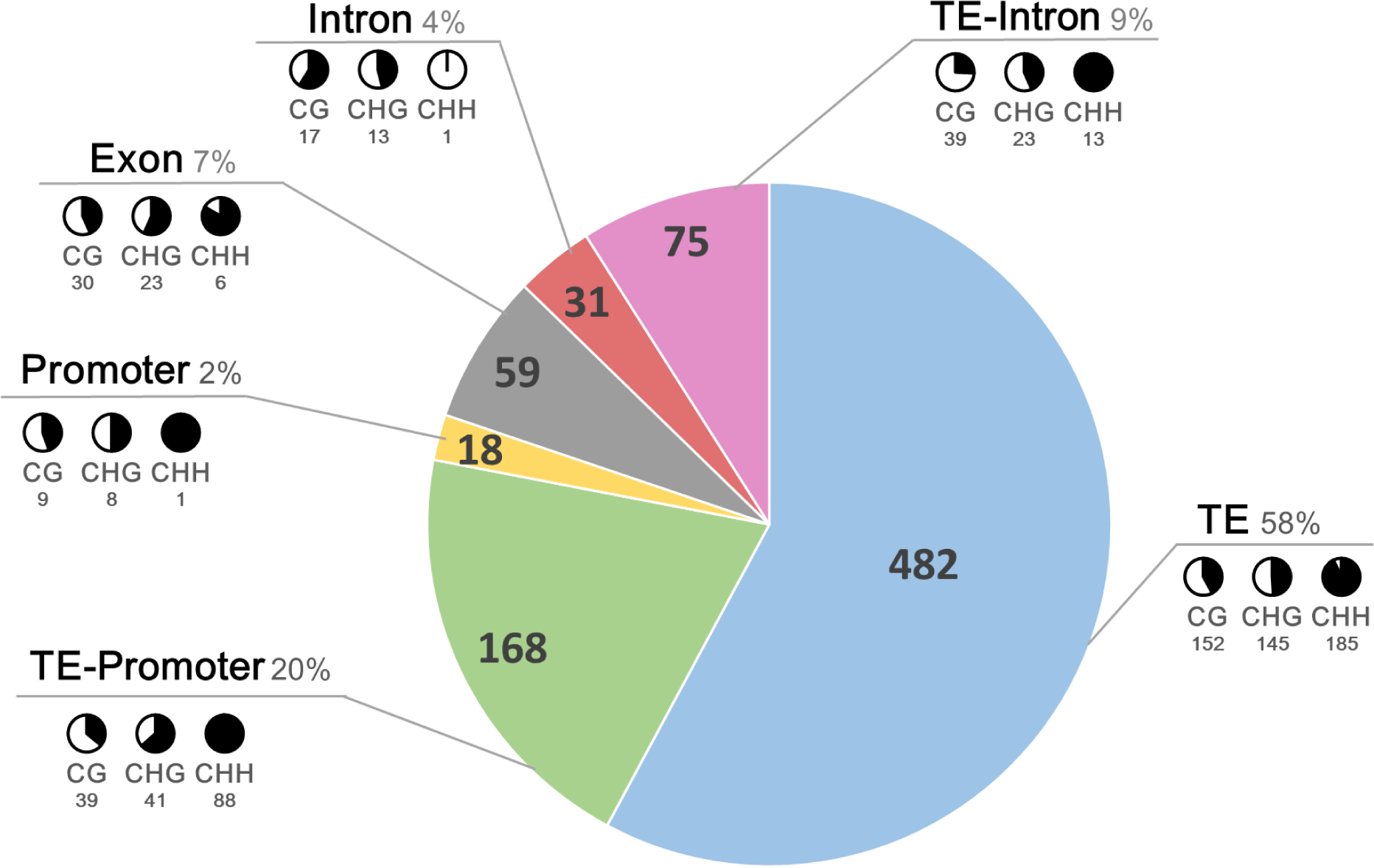
Localisation of identified DMRs in specific genomic features. Black and white pie chart are associated with each cytosine context and represent the proportion of hyper-(black) and hypo-(white) methylated of identified DMRs (number underneath). TE: extragenic Transposable Elements; TE-intron: Intronic Transposable Elements; TE-promoter ; Transposable Element found within promoter sequence.

**Figure 10.**
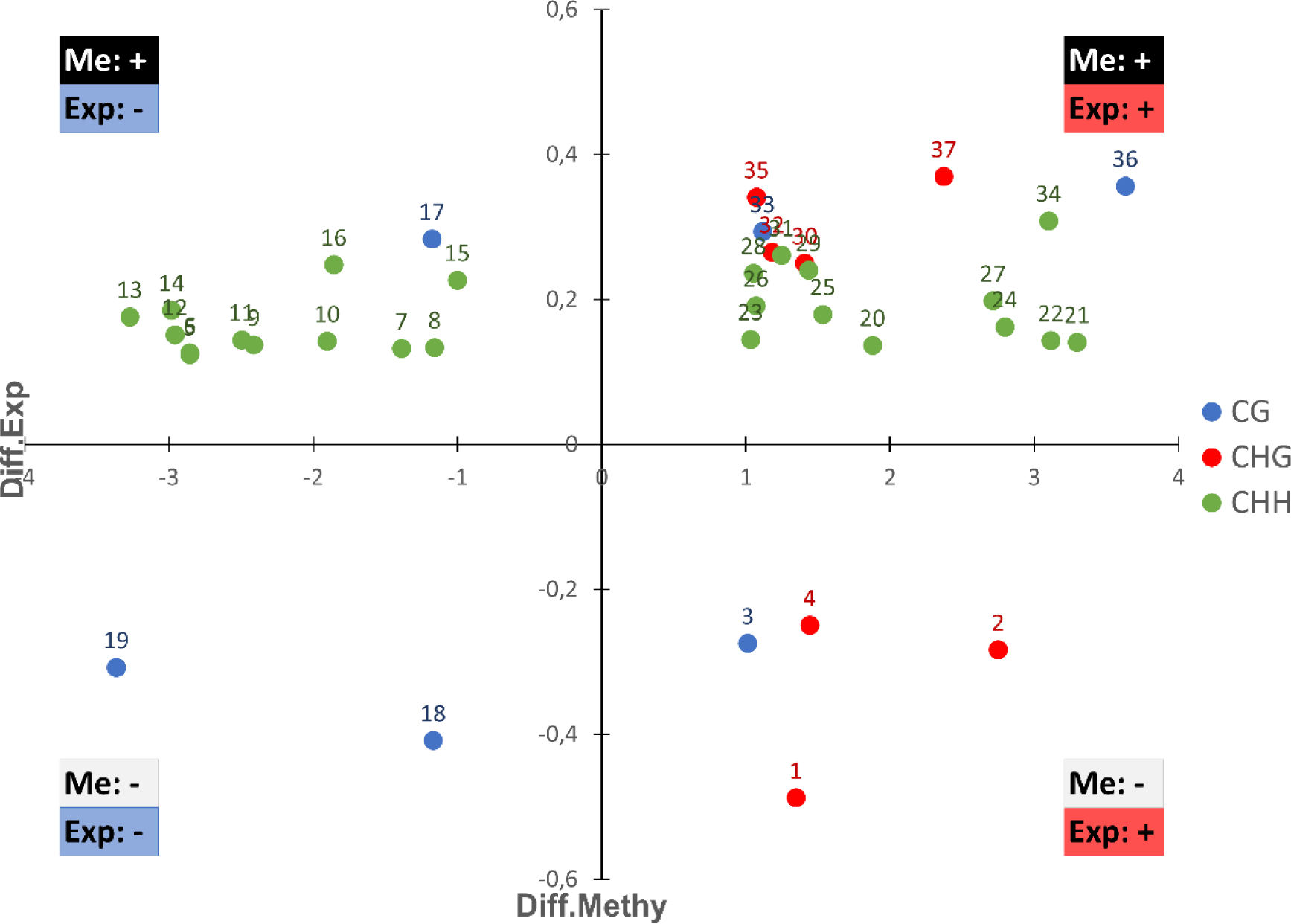
Graphical representation of the relationship between promoter showing differential methylation and associated gene expression changes. Each dot represents a Differentially methylated Region (DMR) found in a Differentially Expressed Gene (DEG) promoter, placed on the graph depending of differential expression value (Log2FC) along the x-axis and its differential methylation along the y-axis. The type of correlation is represented in each corner of the graph. Methylation context is indicated by colors as shown by the legend. Each number represent the N° attributed to the promDMEGs. Me+/-: methylation direction hyper-/hypomethylated; Exp +/-: gene expression up-/downregulated. promDMEGs: Genes with DMRs at promoter showing differential expression.

Overall, sugar limitation modulates the distribution of DNA-methylation genome-wide with a global increase of methylation in grapevine cells. The contexts and types of DMRs seem dependent on their localization (coding or non-coding regions) and of the presence of repetitive elements.

To assess whether genes involved in metabolic pathways showed differential methylation states, the lists of genes containing intragenic-DMRs (exonic, intronic, TE-intronic) or promDMRs (promDMRs, TE-promDMRs) were used for MapMan analysis. Over the 150 intragenic-DMRs identified, 48% (72 over 150) were successfully assigned to a specific function in the considered metabolic pathways (Table S10). These 72 genes belonged to 17 tag categories according to their position within MapMan metabolic pathways (Table S10). Interestingly, 70% (50 over 72) of intragenic-DMRs were associated with DMRs localized in intragenic TE. The same analysis was performed using the genes associated with promDMRs. Approximately 55% (95 over 173) of them could be assigned to a MapMan-referenced metabolic function (Table S11). These 95 genes were classified into 21 tag categories of metabolic functions including the TCA cycle, amino acid, lipid, protein, nucleic acid and hormone metabolism, stress responses, but also transcription regulation and DNA synthesis/repair (Table S11). By contrast, only 3% (3 over 95) of these promDMRs were associated with DMRs localized in TE, suggesting that these genes might be under direct DNA-methylation control.

### Some DMRs are associated with differentially expressed genes

To assess whether changes in DNA methylation might be related with changes in gene expression, we looked for potential overlaps between intragenic-DMRs, promDMRs and DEGs, considering DEGs promoter region as 2kb upstream TSS. In total, 17 intragenic-DMRs showed differential expression (DMEGs; Table S12 and S13), and 34 promDMRs were associated with differential gene expression (promDMEGs). Because the relationship between gene body methylation in transcription regulation remains unclear (Bewick and Schmitz 2017), we focused on the 34 promDMEGs for subsequent analysis (Table 6, Table S14).

**Table 6.**
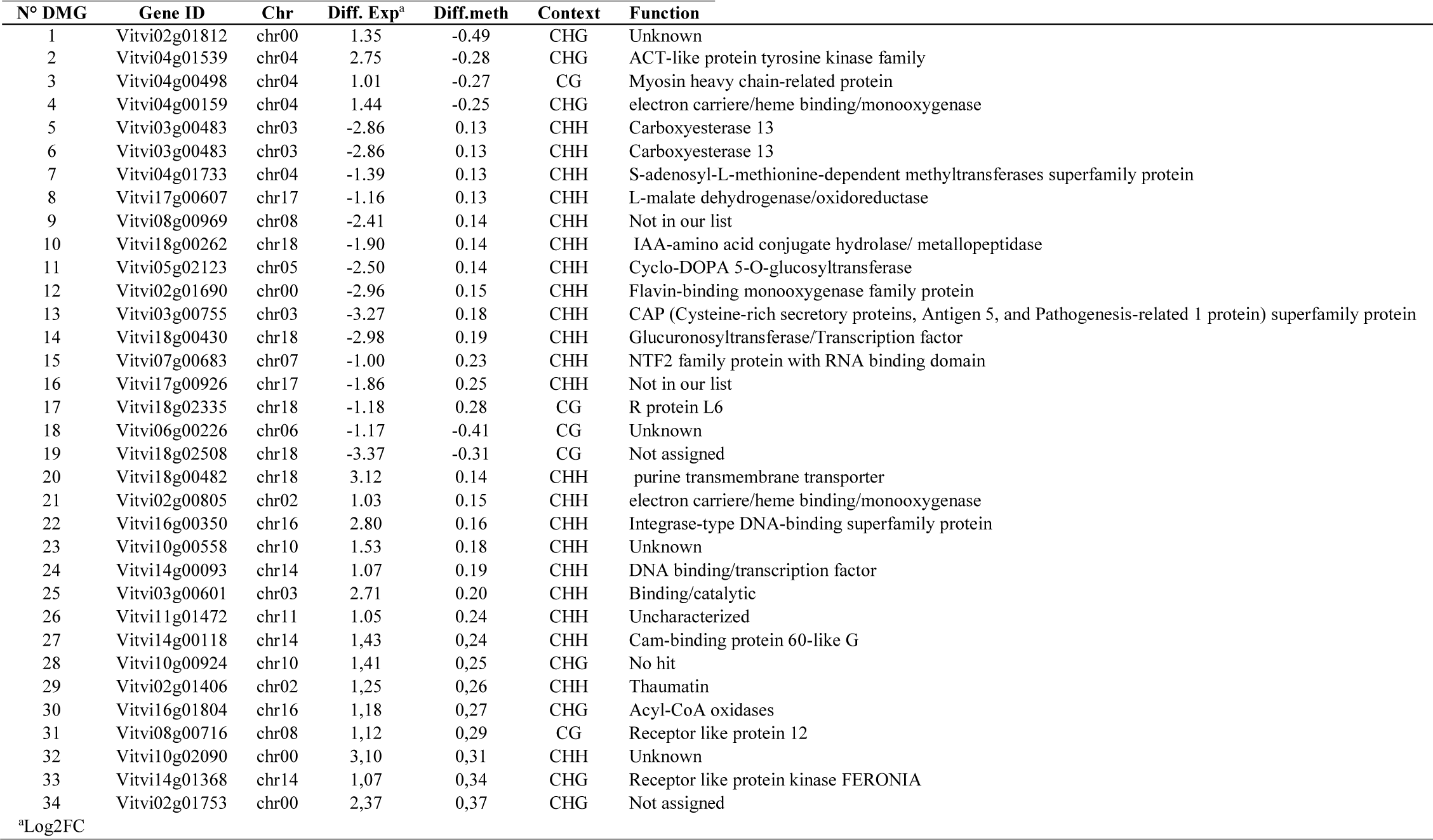
Identified promDMEGs and their associated function. Gene ID: identification based on 12xV2 grapevine genome annotation; Chr: Chromosome; Diff.Exp: Differential expression (Log2FoldChange); Diff.Meth: Difference of methylation between the two conditions; promDMEGs: promDMEGs: Genes with DMRs at promoter showing differential expression.

Consistent with results described above, the majority of the promDMEGs (28 over 34) were hyper-methylated, of which 22 were in the CHH context (Figure 10, Table 6). Among these 34 promDMEGs identified, 24 were associated with genes with an identified putative function (Table 6).

The promDMEGs were further classified in four types based on the relationship between their methylation state (Me: +/-) and differential expression (Exp: +/-; Figure 10). Among the 17 promDMEGs showing negative correlation between methylation state and gene expression (Figure 10, Table 6), genes encoding a SAM-dependent methyltransferase (*Vitvi04g01733*), and a L-malate dehydrogenase/oxidoreductase (*Vitvi17g00607*) showed a hyper-methylated state and reduced expression under sugar depletion (Table 6). Inversely, one gene coding for an electron-carrier coding gene (*Vitvi04g00159*) presented a hypomethylated state related to an increase in expression (Table 6). Interestingly, two promDMEGs also corresponded to genes encoding putative transcription factors (TF), one showing hypomethylated state and enhanced expression (*Vitvi18g00430*) and the other presenting a hypermethylated state associated with enhanced expression (*Vitvi14g00093*). Although the function of these TFs has not been determined in grapevine, *Vitvit14g00093* encode an Homeobox protein of the KNOTED1-LIKE HOMEOBOX (KNOX) transcription factor family (Ensemblplants, UNIPROTKB ref: AC7Q8), orthologous to KNAT6 known for its role in the regulation of meristematic activity in *Arabidopsis* (Belle-Boix et al., 2006), and as such, may be modulating the transcription of specific gene sets (GO:0000981).

Overall, these results suggest that modification of DNA-methylation levels may participate in regulating the expression of different genes in response to carbon limitation.

## DISCUSSION

Carbohydrate depletion results in a coordinated response of grapevine cells at the phenotypic, metabolic, transcriptomic and epigenenomic levels. This was investigated by using heterotrophic CS cells grown in the dark with a 10-fold reduction in glucose concentration (2g/L) as compared to control conditions (20g/L). Because CS cells are completely dependent on external sugar as carbon source, here glucose, they represent an efficient system to control the carbon nutrition of plant material (Morkunas et al., 2012). In this context, our study provides a comprehensive genome-wide analysis of the dynamic of DNA methylation following sugar limitation in plant cells and of their relationships with gene expression and metabolic changes.

### Carbon deprivation leads to cell growth arrest and major metabolic changes in grapevine cells

The transfer of grapevine cells to a medium with reduced glucose content dramatically limits their growth (Figure 1B). This is consistent with previously described data in Arabidopsis and rice cells showing that the response to carbon limitation is rapid (Gout et al., 2011) and influences both metabolisms and phenotypes (Contento, Kim and Bassham 2004; Rose et al., 2006; Wang et al., 2007). More specifically, sugar depletion results in the mobilization of carbon resources, which is characterized by protein degradation, an increase in lipid and starch catabolism, and the recycling of cellular components by autophagy (Contento, Kim and Bassham 2004; Rose et al., 2006; Wang et al., 2007; Morkunas et al., 2012).

This metabolic shift is associated with a profound remodeling of the gene expression patterns 24 to 48h after sugar depletion. In general, sugar-depleted cells show induction of genes involved in nutrient mobilization from starch, sucrose, lipids, as alternative carbon source (Contento, Kim, and Bassham 2004; Wang et al., 2007). Their viability decrease after 48h, and cell death is observed after 72h (Contento, Kim, and Bassham 2004; Wang et al., 2007). Although cell viability was no evaluated in our study, the CS cell biomass stopped increasing and major metabolic changes occurred 24 h to 48 h after subculture. A clear demonstration of these changes was obtained using non-targeted analysis of metabolic profiles, coupled with constraint-based flux analysis. It clearly indicates that most metabolic fluxes are severely limited under carbon depletion, except those involved in the mobilization of cell carbon storage, which were enhanced showing that G- CS cells are experiencing a major metabolic stress.

### Transcriptomic reprogramming is consistent with a major metabolic stress

It is well described that changes in glucose availability to plant cells impact the expression of a large proportion of genes related to stress response, cell wall biogenesis, cellular metabolism reprograming, signaling, and transcription factors (Price et al., 2004). More specifically, end products of photosynthesis such as sucrose and glucose repress photosynthetic gene (Sheen 1990). Of course the carbon-to-nitrogen balance is central to this process (Paul and Pellny 2003). In our conditions, glucose was depleted without modifying nitrogen content, resulting in a reduction of the N/C balance in addition to low glucose availability, which has consequences on plant cell transcriptomic response (Price et al., 2004, Palenchar et al., 2004, Gutiérrez et al., 2007, Huang et al., 2016). Globally these conditions, led to major changes in gene expression profiles (Figure 5). CS cells in G- conditions were characterized by a very strong upregulation of those involved in photosynthesis, particularly in the light reactions (Figure 6B, Figure S8, S10), associated with a strong limitation in biomass production, and the downregulation of genes involved in cell cycle at D6 (Figure 6A). This result is consistent with the study of sucrose-starved non-photosynthetic cells of *Arabidopsis* (Contento, Kim, and Bassham 2004) and demonstrates a strong effect of limiting carbon availability in CS cells.

Consistently, the genes implicated in cell wall synthesis, glycolysis, nucleotides formation and TCA cycle were downregulated, while those involved in metabolic processes associated with carbon mobilization such as the glyoxylate cycle, lipid, starch, and sucrose breakdown pathways was upregulated (Figure 6B, Figure S8, Figure S9, Figure S10). Fluxes involved in the accumulation of resources such as the TCA cycle, and glycolysis also showed a reduced activity in G- versus G+ cells, whereas those involved in resource mobilization including lipid breakdown, cell wall, protein, and starch metabolization were enhanced (Figure S6, Table S5). These results are in line with works showing that after 30-35 h of sugar depletion, plant cells undergo a survival phase, characterized by intensive breakdown of proteins and lipids, hence to a profound modification of cell metabolism (Morkunas et al., 2012), which results in a transient increase in free amino acids. This behavior was suggested to be an adaptive process enabling the cells to maintain a continuous energy supply under sugar starvation (Morkunas et al., 2012).

### THF and SAM cycle are downregulated in grapevine cells under carbon limitation

Among the different metabolic pathways, we have focused on the 1C metabolism, as it is critical for all cell transmethylases by providing SAM, the universal methyl donor. More specifically, the synthesis of methylation precursors is dependent on the activity of the THF and SAM cycles (Lindermayr et al., 2020). At D6, transcriptomic and fluxomic analyses demonstrate both downregulation of genes involved in methylene-THF, and a reduction of fluxes of the tetrahydrofolate pathway in G- conditions (Figure 4 and 6, Figure S10, Table S6). However, fluxes implicated in dihydrofolate (DHF) synthesis where not off under sugar limitation (Figure 4, Table S6). Apart from the fact that DHF are THF precursors (Hanson and Gregory 2002, 2011), and that DHF conversion to THF might be related to redox balance maintenance to cope with oxidative stress through NADPH production (Gorelova et al., 2017) their role in the cell remains unclear.

Overall, grapevine cells facing glucose deficiency are engaged in a survival phase that contributes to maintaining their primary functions by using intracellular carbon resources available. This results in a profound modification of the cell metabolic fluxes that also the 1C metabolism. As the 1C metabolism includes tetrahydrofolate synthesis (Jabrin et al., 2003) and SAM cycle (Moffatt and Weretilnyk 2001), sugar depletion may result in a limited availability of SAM, the universal methyl donor.

### Sugar depletion impacts the expression of genes associated with DNA-methylation regulation

We identified four differentially expressed grapevine DNA methyltransferases (*VvMET1*, *VvCMT1*, *VvCMT3*, *VvCMT4*), one helicase (*VvDDM1*) and one demethylases (*VvDML3*) genes in G- condition (Table 2), suggesting that DNA methylation process was affected at D6 in grapevine cells under carbon limitation. The DDM1 protein is a chromatin remodeling factor required for the maintenance of CG methylation (Vongs et al., 1993) while VvMETs and VvCMTs are DNA methyltransferases implicated in the CG and the CHG methylation maintenance process respectively (Cao et al., 2003). *VvMET1* was let aside for further analysis as it showed very limited changes in expression level between G- and G+ conditions (0.45 L2FC, padj=0.002). The *VvDDM1* and *VvCMTs* genes were all repressed under carbon limitation (from -1.06 to -1.45 Log2FC), which is consistent with a reduced need for DNA methylation maintenance processes as G- cells undergo cell division arrest. *VvDML3* is also repressed in G- condition (-2.60 L2FC) as compared to control, suggesting a reduction of the active demethylation process as well (Table 2).

### Carbon limitation induces global DNA methylation changes

Plant cell DNA-methylation landscape is modified in response to environmental changes in plants, including nutrient starvation such as phosphate limitation (Akhter et al., 2021; Yong-Villalobos et al., 2015; Tian et al., 2021). Here, we have shown that global cytosine methylation is higher in G- than in G+ condition with a 4%, 8% and 20% increase in the CG, CHG and CHH context respectively (Figure 7, Table 3, Table 4). Consistently, the DNA-methylation level within genes and in 2 kb regions up/downstream of genes is higher in the CHG and CHH contexts in G- as compared to G+ conditions (Figure 7B). A similar observation was made for TE (Figure 7B). Such results are consistent with recent studies showing that mCG is rapidly maintained, while the reestablishment of mCHG methylation is delayed leading to asymmetrical methylation throughout the cell cycle, and mCHH is depleted in actively dividing cells (Borges et al., 2021).

In our conditions, at D6, G+ cells are dividing actively (Figure 1B), while G- cells have most likely stopped dividing (Figure 1B), considering the strong downregulation of cell cycle-related genes (Figure 6A). Thus, the hypomethylation observed in G+ versus G- cells might in part reflect a difference in cell division activity between the two conditions.

However, we have also shown that four ARGONAUTE encoding genes (*VvAGO10a, VvAGO11, VvAGO5 and VvAGO2a*) showed higher expression levels in G- cells (Table 2). Although the functional characterization of these proteins in grapevine is lacking, they are orthologous to genes involved in the RdDM in *Arabidopsis* (Chapman and Carrington 2007). As such they are part of the complex machinery leading to the synthesis of siRNAs and thereby to the CHH methylation through the RdDM pathway (Erdmann and Picard 2020). Similarly, the *VvAGO, VvDCL* and *VvRDR* gene families were shown to be under complex regulations in plants under abiotic stress such as water deficit and salt treatment (Zhao et al., 2015). Indeed several works have shown that CHH methylation is highly dynamic in plants submitted to stresses (López et al., 2022; Lin et al., 2022). This suggests that the RdDM also contributes to the hypermethylation at CHH sites under carbon limitation, contributing to changes of the methylation landscape.

### The majority of DMRs are localized in transposable elements

We further investigated DNA-methylation by determining the differentially methylated regions between G- and G+ cells 2 days after subculture. The 848 DMRs identified were equivalently distributed between the three C sequence contexts CG, CHG and CHH, which indicates that methylation was dynamically changed after 48h exposure to carbon limitation (Figure 8). While a near equal number of hypo- and hyper-DMRs were identified in CG and CHG context, CHH showed a strong bias toward hyper-DMRs under sugar limitation (Figure 8). These DNA-methylation changes mostly concern TE region, irrespective of C sequence context and genomic locations (Figure 9), suggesting that TEs methylation changes are more likely to occur in G- cells.

The DMRs located in intergenic-TEs and Promoter-TEs showed an overrepresentation of hyper-DMRs in the CHH context, as compared to CG and CHG context for which we observed equivalent repartition between hyper- and hypo-DMRs (Figure 9). Similar profile of CHH hyper-methylation has been recently reported in tomato under phosphate starvation (Tian et al., 2021). The majority of identified DMRs were localized at TE, although some of them were found within intronic regions and gene promoters (Figure 9). We identified 165 DMRs within genes (Figure 9) among which 75 are located at intronic-TE. Our results are in accordance with literature where hyper-methylation of TEs across all chromosomes in response to stress has also been recently reported for the first time in maize (Achour et al., 2019), as well as in tomato suffering from phosphate starvation where CHH DMRs were enriched in intra/inter-genic TE (Tian et al., 2021). As TEs are targeted by repressive epigenetic marks as a preservation mechanism for genome integrity (Okamoto and Hirochika 2001), this response has been proposed to be related to the activity of the RdDM pathway in repressing transposons in accessible chromatin environments (Gent et al., 2014).

Overall, our results show that sugar limitation results in important changes in DNA methylation patterns in all sequence contexts that mostly concern TE sequences. Furthermore, the high number of hypermethylated cytosine in the CHH argues in favor of the involvement of RdDM in response to sugar limitation.

### Differential methylation may have an impact on gene expression in cells under carbon limitation

Among the 848 DMRs identified, 165 (19%) are located in genes and 186 (22%) in their promoter region (-2kb upstream the TSS; Figure 9). In both cases, DMRs were enriched in genes related to the central and secondary metabolism, along with those involved in stress response, photosynthesis, chromatin structure, protein PTMs, and transcription regulation (Table S10). Interestingly, the category of genes containing DMRs is similar to those of DEGs. Among these DEGs, 51 over the 5,607 identified also presented DMRs in their promoter regions among which 17 presented a negative correlation between the methylation state and gene expression. These 17 promDMEGs include genes coding for putative transcription factor such as Class 1 KNOX gene, enzymes involved in the central metabolic pathways including a malate dehydrogenase, and an S-adenosyl-methionine-dependent methyltransferase among others (Table 6). This observation strongly suggests that DNA-methylation is involved in the control of gene expression as an integrated part of the cell metabolic and molecular responses to carbon depletion. Similarly, differentially expressed genes related to phosphate starvation regulation in tomato plants under low phosphate conditions display DNA-methylation changes (Tian et al., 2021). Hence, DNA-methylation dynamic may represent a general strategy of plants cells to deal with nutritional stresses, including carbon depletion.

### Carbon limitation could impacts other epigenetic mechanisms

As for DNA-methylation, Histone Post Translational Modifications (HPTMs) also rely on the availability of metabolic precursors and cofactors. Histone methylation depends on SAM availability. Studies with Arabidopsis uncovered complex interactions between DNA and histone methylation, showing that DNA-methylation was tightly correlated with an enrichment in H3K9me2 marks (Du et al., 2015). Similar to the inhibition of DNA methyltransferase encoding genes described above, all differentially expressed genes encoding histone methyltransferases identified were downregulated in cells undergoing carbon depletion (Table 2). This suggests a global inhibition of most if not all methylation processes associated with epigenetic modifications in G- cells. Furthermore, the TCA cycle was inhibited in our cells. In mammals, experiments leading to disruption of TCA-cycle carbohydrate fluxes influenced 2-OG production, resulting in subsequent modification of DNA and histone methylation profiles (M. Xiao et al., 2012). Although the exact mechanism was not identified it was suggested that JMJC proteins 2-OG dependent histone demethylases could be involved.

Inhibition of fluxes involved in TCA cycle was observed in G- cells that may lead to similar consequences eventually involving the JMJC orthologues identified in grapevine. In addition, the down-regulation of genes encoding the putative grapevine JMJC in G- cells (Table 2) further indicated that they could participate to a potential metabolic effect that would result in histone hypermethylation.

Overall, these results show that epigenetic changes under carbon shortage are unlikely to rely exclusively on DNA-methylation. The differential expression of histone modifiers in G- versus G+ cells suggests that the distribution of histone marks may also be affected under sugar depletion. Further analysis is now needed to assess the distribution of HPTMs and their relation with gene expression.

In summary, our study provides a comprehensive genome-wide analysis of DNA-methylation changes and their direct relationship with gene expression to orchestrate metabolic adjustment in the response of plant cells to sugar limitation. This approach aimed at understanding the regulatory networks, which coordinate plant cell adaptation to carbon deficiency and the potential role of epigenetic mechanisms in this process. As shown in this work depletion of carbohydrates has a profound impact on the metabolism of cells and leads to an important remodeling of DNA-methylation landscape and gene expression patterns.

In particular, we have demonstrated an important DNA-methylation remodeling under carbon depletion, that was associated with changes in gene expression in a few cases. Genes that were targeted include genes encoding enzymes involved in the metabolism of cells, but also transcription factors that in turn may coordinate the response of cells to carbon depletion. Hence DNA-methylation, in association with other epigenetic mechanisms, could contribute to plant cell responses to carbon depletion.

## MATERIALS AND METHODS

### Cell culture

Grape cell suspensions of Cabernet Sauvignon Berries (CSB, Atanassova et al., 2003) initiated from fruit tissues were cultured in erlenmeyer of 250 mL containing 50 mL of a modified liquid Murashige and Skoog medium (MS, Sygma ref M0221), supplemented with vitamins (0.025 g/L of *myo-*inositol, 0.25 mg/L of nicotinic acid, calcium pantothenate, HCL pyridoxine, HCL thiamine and 0.0025 mg/L of D(+)-Biotin), 0.6 mg/L of benzylaminopurine (BAP), 2.3 mg/L of naphthalene acetic acid (NAA), 0.25g/L of casein enzymatic hydrolysate (Sigma C0626), and 20g/L of glucose and were maintained at 23°C in darkness on an orbital shaker (120 rpm). Cells were subcultured weekly by transferring 10 mL of the cell suspension to 40 mL of fresh medium, in individual vials for each time point and condition, thereby avoiding any opening stress before sample collection. Sample collection was performed every day from the day of inoculation (D0) to D10 at the same hour. Cell growth was estimated by measuring the fresh weight (FW) of cells contained in 30ml of the cell suspension after removal of the culture medium by filtration. Cells and culture medium were subsequently frozen in liquid nitrogen, ground in this state to a fine powder in a Retsch Mixer Mill MM 400 (Fischer Scientific), at 30 Hz for 1 min, and stored at -80°C until processed.

To evaluate the consequences of a limitation in sugar availability, cells were cultured during four days in standard condition (SD) and subsequently transferred to either a glucose rich (G+: 110 mM glucose (20g/L)) or glucose poor (G-: 11 mM glucose (2g/L)) medium. To achieve this, cells were decanted for a few minutes, and the buffer removed by pipetting. Cells were washed three times with either G+ or G- fresh medium, depending on the conditions, and re-suspended in the appropriate medium. This led to a moderate loss of cells, roughly ∼40 out of ∼195 gFW/L after subculture in either type of medium (Figure S1). Cells were grown in these new conditions for 6 additional days. The cell water content was evaluated using ∼140 to 200mg FW of frozen (maintained at -80°C) grounded cells, by comparing the cell mass before and after 36h of lyophilization.

### Metabolic analysis

#### Soluble sugars content in culture medium

Glucose and fructose concentrations were measured enzymatically with an automated absorbance microplate reader (Elx800UV, Biotek Instruments Inc., Winooski, VT, USA) using the glucose/fructose kit from BioSenTec (Toulouse, France) according to manufacturer’s instructions. The results were presented as ‘extracellular glucose’ concentration.

#### Metabolite extraction

Extractions for both targeted and untargeted metabolite analyses were performed in quadruplicate for each sample of frozen cell powder at -80°C. Extraction was essentially performed as described in Luna et al., (2020) with 4 biological replicates (n=4) at the *Bordeaux Metabolome Facility* (https://metabolome.u-bordeaux.fr/en/, Villenave d’Ornon, France) with the following slight modifications: (1) 20 mg of fresh frozen-powder (FW) of each replicate sample were used for extraction, (2) ∼10 mg of poly(vinylpolypyrrolidone) (PVPP) were added in each tube prior to extraction to precipitate polyphenols.

#### Targeted metabolite phenotyping

Targeted analyses of soluble sugars (glucose, fructose, sucrose), starch, malate, citrate, total soluble proteins, total amino acids were conducted at *Bordeaux Metabolome Facility*. Measurements were based on coupled enzyme assays as described previously by Biais and coworkers Biais et al. (2014), except for total soluble proteins measured by Bradford assay (Bradford 1976).

#### Untargeted Metabolic Profiling

Untargeted metabolic analysis were performed using UHPLC-LTQ-Orbitrap mass spectrometry (LCMS) with an Ultimate 3000 ultra-high-pressure liquid chromatography (UHPLC) system coupled to an LTQ-Orbitrap Elite mass spectrometer interfaced with and electrospray ionization source (ESI: ThermoScientific, Bremen, Germany), following the same method as described Dussarrat et al., 2022; 2023. Each experiment was performed using 4 independent biological replicates (n=4). The analytical sequence consisted of 81 samples, 9 extraction blanks (prepared without biological material and used to rule out potential contaminants detected by untargeted metabolomics), and 10 Quality Control (QC) samples that were prepared by mixing 20 µL from each sample. QC samples were used for i) the correction of signal drift during the analytical batch, and ii) the calculation of coefficients of variation for each metabolomic feature so only the most robust ones are retained for chemometrics (Broadhurst et al., 2018). Raw LCMS data were processed using MS-DIAL software (v. 4.7; Tsugawa et al., 2015) following optimised parameters (see Supplemental Material 1) and yielded 8 881 features. Putative annotation of differentially expressed metabolites was performed by exploiting the MSDIAL online library MSMS-Public-Neg-VS15.msp (36,848 records) and screening of the MS1 detected exact HR *m/z* and MS2 fragmentation patterns against multiple online databases (http://prime.psc.riken.jp/compms/msdial/main.html#MSP; Tsugawa *et al*., 2015). After data-cleaning (blank check, SN > 10, CV QC < 30%), 1719 metabolomic variables were retained for further chemometrics. The final dataset was normalized (median normalization, cube-root transformation and Pareto scaling) using MetaboAnalyst (v 5.0; Pang *et al*., 2021) prior to multivariate statistical analyses. Each experiment was performed using 4 independent biological replicates (n=4)

#### Amino acids quantification by HPLC

Samples extracts were derived with 6-aminoquinolyl-*N*-hydroxysuccinimidyl carbamate (AccQ-Tag derivatization reagent, Waters, Milford, MA, USA) according to Arrizabalaga-Arriazu et al. (2020) and the references within, using an UltiMate 3000 UHPLC system (Thermo Electron SAS, Waltham, MA USA) equipped with an FLD-3000 Fluorescence Detector (Thermo Electron SAS, Waltham, MA, USA). Free amino acids’ separation was achieved by using an AccQ-Tag Ultra column, 2.1 × 100 mm, 1.7 μm (Waters, Milford, MA, USA) at 37 °C with elution at 0.5 mL/min (eluent A, sodium acetate buffer, 140 mM at pH 5.7; eluent B: acetonitrile; eluent C: water). Chromatographic analyses were carried out using an excitation wavelength of 250 nm and an emission wavelength of 395 nm. The identification and quantification of 19 amino acids (excluding tryptophan and cysteine) were done as previously described by Arrizabalaga-Arriazu et al. (2020).

#### Quantification of NAD(P) contents in grapevine cells

Extraction of NAD+, NADH, NADP+ and NADPH was performed from 20 mg of fresh frozen grapevine cell powder (FW) with the addition of ∼10mg of PVPP prior to extraction and quantified as described by (Decros et al., 2019; 2023). Quantification was performed in technical duplicate using four independent biological replicates.

#### Cell wall content estimations

Analysis of cell wall biomass performed on the HitMe plateform at *Bordeaux Metabolome Facility* from 20 mg of each replicated sample following plate preparation and randomization described above, following the platform protocol. Briefly, after metabolite extraction, pellets were separated from supernatant and resuspended in 250 µL of NaOH 0.5 M at 95°C for 20 min. Supernatant was removed after centrifugation, the pellet was washed twice by addition of 250 µL of water and centrifuged at 2500 rpm for 10 min. After one night of lyophilization, cell wall content was estimated by calculating the difference of tube weight before and after discarding the pellet.

### Flux-Balance Model

The flux-balance model of heterotrophic plant central metabolism was initially described in Colombié et al., (2015) and improved (details given in Lacrampe et al., *under* review) to better describe all the reactions involving energy, metabolic compounds for biomass synthesis (proteins, cell wall, lipids, nucleotides, amino acids) and metabolic precursors for secondary metabolism. Briefly, this model (Tables S1-S2, Supplementary Material 2) took into account the main pathways, such as glycolysis, tricarboxylic acid (TCA) cycle, oxidative pentose phosphate pathway, sucrose catabolism, etc, using glucose as carbon source (*Vglc_up*), nitrate as nitrogen source (*Vno3_up)* and glutamine as both (*Vgln_up*). All the cofactors were defined as internal metabolites, which means that they were balanced, thus constraining the metabolic network not only through the carbon and nitrogen balance but also through the redox and energy status.

The model described the cell metabolism through a set of 302 reactions involving 177 internal metabolites. At steady state, the mass balance equation was expressed by

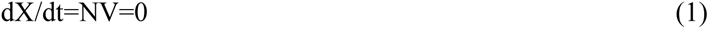

X is the vector of internal metabolites, V the flux vector composed of n reactions of the network, and N the stoichiometry matrix. To solve the system, constraints were applied on each flux. First, thermodynamic properties were used to constrain fluxes from reversibility to irreversibility. Thus, among the internal reactions of the metabolic network, lower bounds of irreversible reactions were set to zero. The most important constraints were the external fluxes, also called exchange fluxes. They were calculated from the accumulated metabolites and biomass components, covering an average of 75% of the dry biomass under control and carbon-limiting conditions (Table S3). Similarly, the flux of glucose uptake rate was calculated from extracellular glucose concentration (g/L). To achieve this, the best polynomial fitting (of experimental data expressed in mmol/L) was searched (criteria r2 max) and the corresponding flux was calculated by derivative (Table S4). Finally, flux minimization, which leads to a unique solution (Holzhutter, 2004), was used as the objective function to solve the system, and generate flux maps each day before and after carbon limitation (Table S5).

#### Software

Stoichiometric model (in *sbml* format, supplementary material) and mathematical problems were implemented using MATLAB (Mathworks R2018, Natick, MA, USA), solver *quadprog* with interior-point-convex algorithm for the minimization.

### Molecular characterisation

#### Nucleic acid extraction

Both DNA and RNA were both extracted from the same sample (∼200mg cells of resh frozen powder (FW)) of independent biological replicates (n=3) of each conditions (D4, D6 G+, D6 G-) according to (Berger et al., 2022).

#### Bioinformatic analysis of RNAseq data

High-throughput sequencing of RNA samples were performed using DNBSEQ Sequencing technology (pair-end, 150bp) service provided by BGI-Genomics platform (http://bgi.com). We generated 9 cDNA libraries corresponding to three biological replicates for each group (D6_G+ and D6_G-). Raw reads were trimmed using Trimmomatic v0.38 in PE mode (Bolger et al., 2014), The sequence alignment files were generated by STAR (version 2.5.1b) (Dobin et al., 2013). To generate the raw gene counts, we used the feature Count function of the Rsubread package (Liao, Smyth, and Shi 2014) which assigns mapped sequencing reads to genomic features based on the grapevine reference genome assembly of PN40024 12X.2 (https://urgi.versailles.inra.fr/Species/Vitis/Data-Sequences/Genome-sequences). We used the DESeq2 package to identify differentially expressed genes (Love, Huber, and Anders 2014). After selection of genes of which ⅓ of the values present a CPM (count per million reads) >10 for ⅓ of the minimum values, differentially expressed genes (DEGs) were identified using a False Discovery Rate (FDR)-adjusted p-value threshold < 0.05. In addition, only DEGs with a log2 fold change > 1.0 were selected.

#### Bioinformatic analysis of WGBS data

Whole Genome Bisulfite Sequencing of DNA samples were performed using DNBSEQ-sequencing (pair-end, 100bp) technology provided by BGI-Genomics. Reads obtained from the WGBS approach were first trimmed with TrimGalore! (Version v0.4.5). First, we assessed the bisulfite conversion rate using the unmethylated grapevine chloroplast genome with Bismark tool (version v0.20.0) (Krueger and Andrews, 2011). Cleaned reads were then aligned onto the grapevine reference genome PN40024 (12Xv2) using Bismark (version v0.20.0) allowing 5 mismatches. Reads with multiple alignment were discarded. PCR duplicates were also removed using the ‘deduplicate_bismark’ tool. The methylation state of each cytosine was calculated, for the three contexts CG, CHG and CHH. We then used the DSS R package (dispersion shrinkage for sequencing data - version 2.39.0) (Wu et al., 2015) to identify differentially methylated regions (DMRs) based on a Wald test procedure and accounting for both biological variations among replicates and sequencing depths. First, differential methylation statistical tests were performed at each C locus by calling the DSS DMLtest function with the parameter smoothing=TRUE. Then, differentially methylated loci were retained when the difference in mean methylation levels was >0.1 for CG or CHG contexts and >0.07 for CHH context with a posterior probability >0.9999. DMRs were then identified using the DSS callDMR function with standard parameters. To define hypo- or hyper-DMRs, we applied a cut-off of at least a 10%, 25% and 25% change in methylation ratio for CHHs, CHGs, and CGs, respectively.

#### MapMan analysis

All the gene models were automatically categorized according to the MapMan ontology (x3.6) with Mercator tool (Lohse et al., 2014) and MapMan standalone software v3.5.1 (Thimm et al., 2004) was used to explore the data.

### Statistical analysis

Untargeted metabolic profiling data were checked for statistical significance by ANOVA for global variation. Significant differentially accumulated metabolites among conditions considered (Fisher’s P<0.01) were determined and analyzed using MetaboAnalyst 5.0 online software (https://www.metaboanalyst.ca/docs/Format.xhtml).

For all other analyses, mean and confidence interval values (CI) were calculated with R software (R-studio, R 4.1.1). Data were checked for normality using Shapiro-Wilk test, and further checked for significance by performing parametric (t-test) or non-parametric (Wilcox-Mann-Whitney test) depending on their normality status.

## Supporting information

MSDIAL parameters

Stoichiometric Model

## Aknowledgments

We are grateful to the genotoul bioinformatics platform Toulouse Occitanie (Bioinfo Genotoul, https://doi.org/10.15454/1.5572369328961167E12) for providing help, computing and storage resources.

