## Supplementary Figures for "Grapevine cell response to carbon deficiency requires transcriptome and methylome reprogramming"

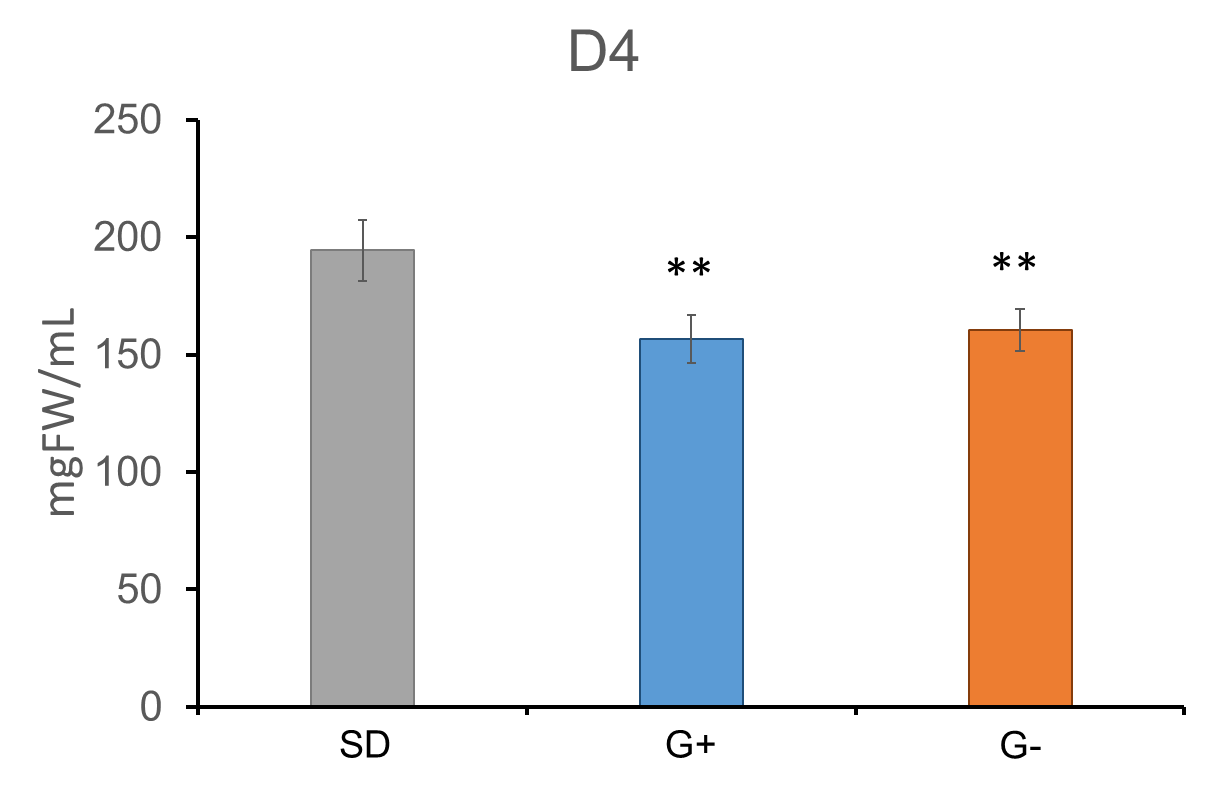

**Figure S1: Fresh weight (FW) after medium change between Standard (SD) and G+/- conditions. Cell concentration presented in mgFW/mL was estimated before (SD) and after the transfert in a new medium (G+ and G-). **: pval ≤ 0.01**

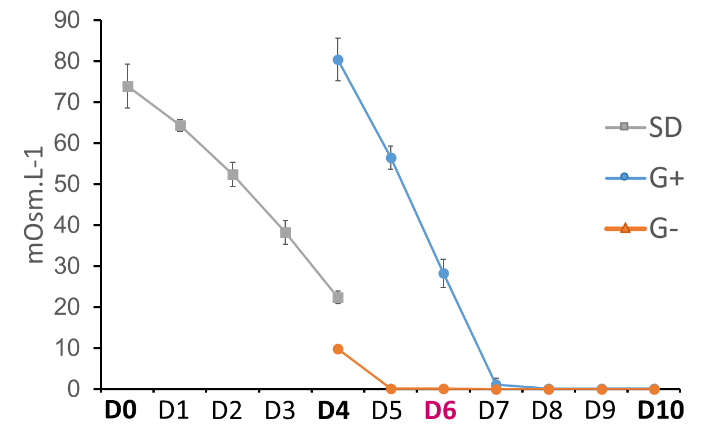

**Figure S2: Medium osmolarity evolution of SD, G+ and G- conditions from D0 to D10.**

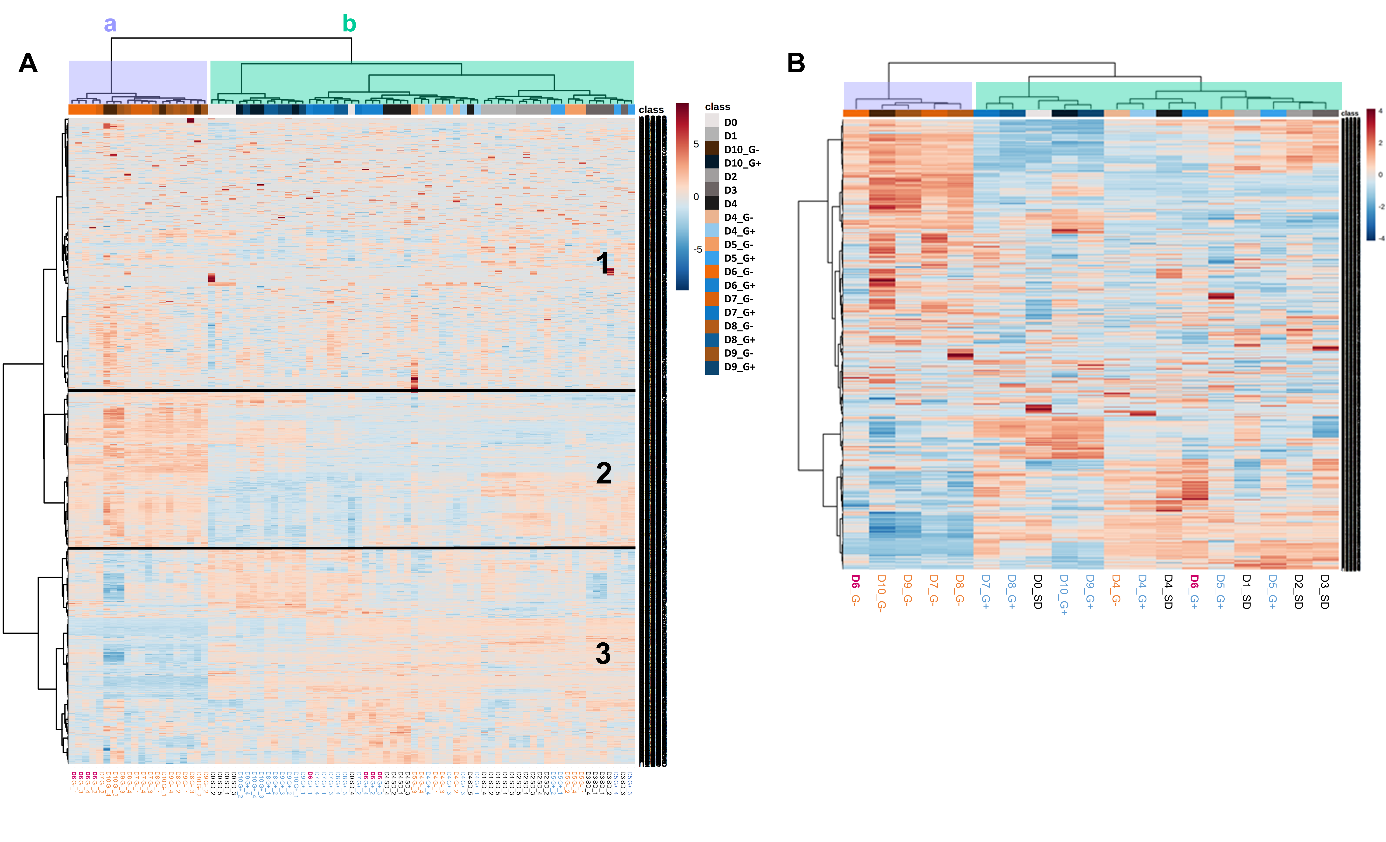

**Figure S3: Heatmap representation of samples metabolic profiling based on the enrichment value of the 1719 metabolic signature (normalized per row) detected using LCMS analysis, and clustered using Pearson’s correlation. (A) Representation of metabolic profile of each sample replicates ate each time point (n ≥ 4) and (B) averaged metabolic profile per day and condition.** The heatmap presented in (A) was separated in three horizontal parts (1, 2, 3) according to the pattern of metabolic features of accumulation within the samples. Part 1 groups the metabolites that are not differentially accumulated depending on the samples. The second group (2) represents metabolites that are more accumulated in D6 to D10 G- samples, and the last one (3) is composed of metabolites less detected in D6 to D10 G- samples, showing the clear differentiation between G+ and G- condition from D6, the latter illustrated by a vertical separation. Samples separated in two cluster (a, light purple; b, light green) based on their metabolic features.

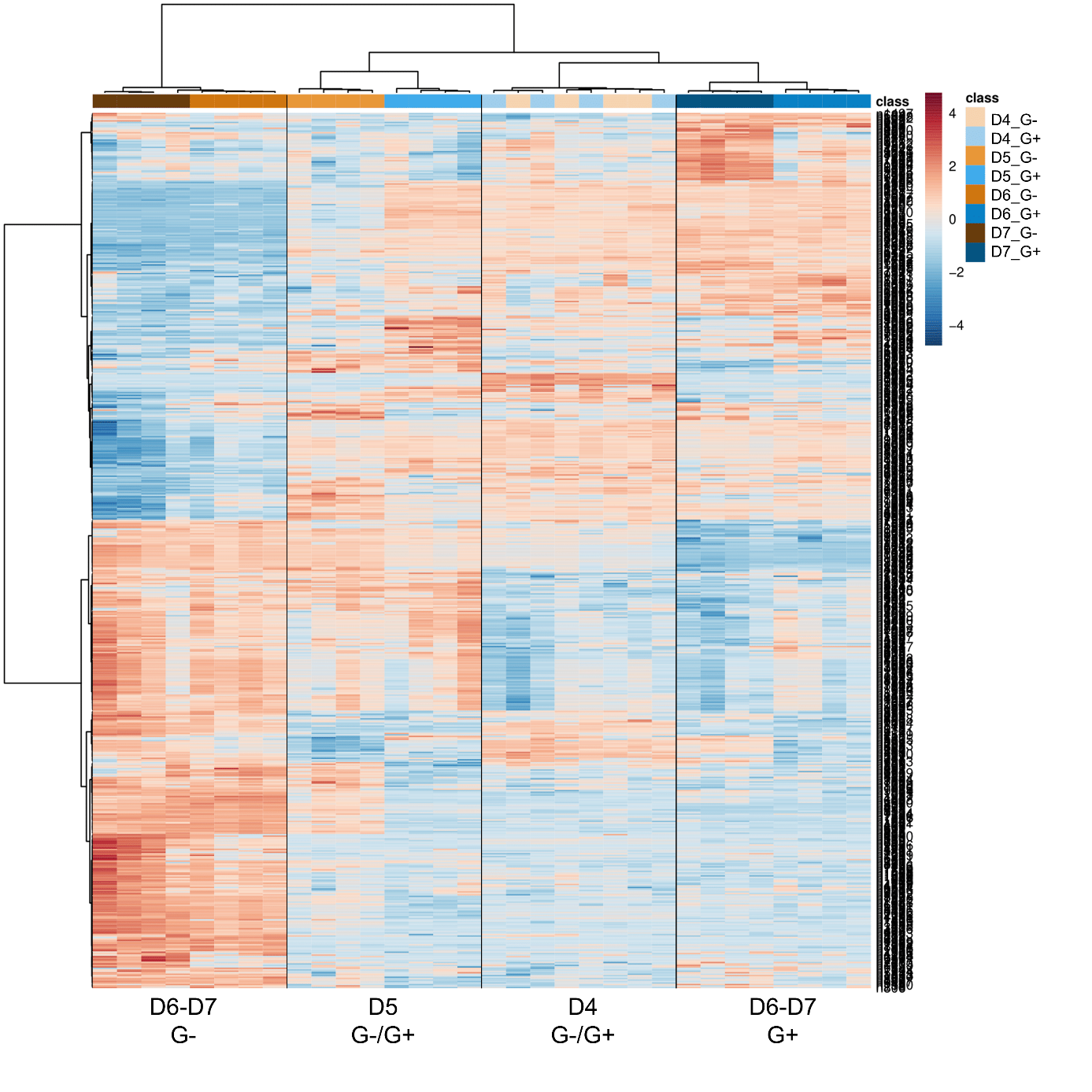

**Figure S4: Heatmap representation of samples profiles from D4 to D7 in G+ and G- conditions based on the 596 most significantly differently accumulated metabolic features across all conditions (p < 0.01)**

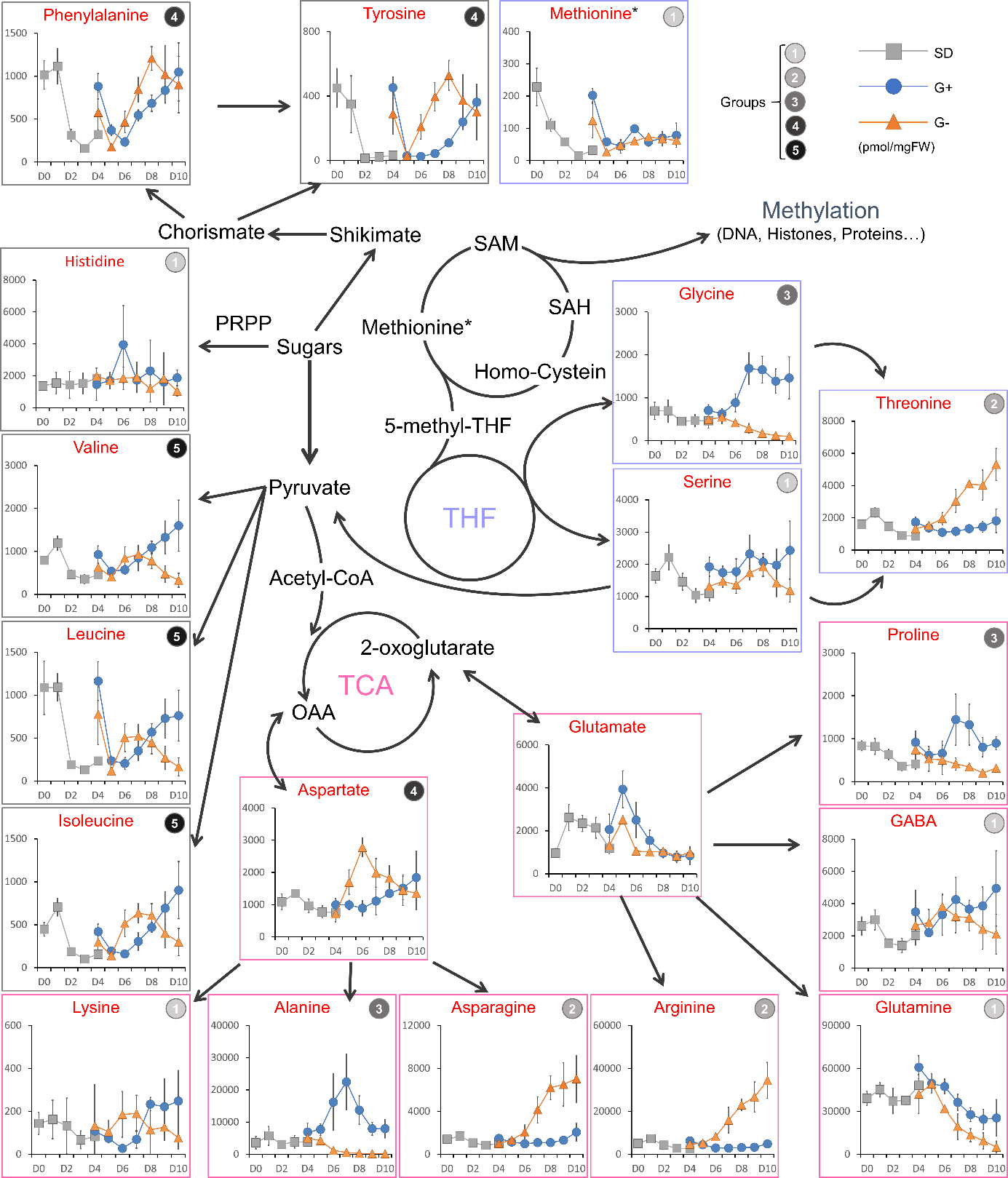

**Figure S5: Amino-acids accumulation (y-axis, pmol/mgFW) in grapevine cells harvested from D0 to D10 (x-axis) in standard (SD, grey), G+ (blue) and G- (orange) conditions. Amino acids pathways are inferred according to the Kyoto Encyclopedia of Genes and Genomes (**[**https://www.genome.jp/kegg**](https://www.genome.jp/kegg)**) and published work about THF and Methionin cycle (Meng et al., 2018, doi: 10.1104/pp.18.00183). Vertical bars indicate CI (n=4).**

According to their accumulation profiles, we identified five types of variations in amino acid abundance. (1) Methionine, glutamine, lysine, GABA, serine and histidine did not show major differences in accumulation pattern between G+ and G- conditions from D4 to D10 although glutamine contents in G+ were slightly higher than in G- cells. The second group (2) gather asparagine, arginine and threonine, showing contrasted profiles between G+ and G- conditions. Indeed while no accumulation variation was detected in G+ condition, G- cells showed an exponential increase of these compounds from D4 to D10. By contrast, the third type of accumulation profile (3) concerns alanine, proline, and glycine contents, characterized by a progressive decrease from D4 to D10 in G- cells, while G+ cells displayed a strong increase from D4 to D7 followed either by a plateau, or a slight decrease from D8 to D10. The fourth group (4) is composed by phenylalanine, tyrosine, and aspartate. Except for aspartate, these compounds showed similar accumulation profiles in G+ and G- condition at D4 and D5. In G+ cells, the three amino acids were progressively accumulated from D6 to D10. In G- cells, these compounds are strongly accumulated and form peak at D6 (aspartate) or D8 (phenylalanine and tyrosine) followed by a decrease until D10. The last group (5) include valine, leucine and isoleucine characterize by a decrease from D4 to D5 followed by exponential accumulation in G+ cells from D6 to D10. All three amino acids are rapidly accumulated in G- cells to reach a plateau at D6, followed by a progressive decrease from D8 to D10. Glutamate was not assigned to a group according to its singular behavior. In G+ cells, glutamate contents increased to a peak (D5), and progressively decreased to stabilize at D8 at value observed at D4. In G- cells, equivalent and sequential increase and decrease are observed between D4 and D6 to stabilize at the same value as the one observed in G+ condition

**Figure S6.: Metabolic reorganization between G+ and G- cells. The flux map shows the differences in simulated fluxes between G- and G+ with increased fluxes in G- cells represented in orange, increased fluxes in G+ cells represented in blue, and unchanged fluxes in grey.** Fold-changes in flux intensities between G- and G+ were calculated as abs(log(abs(G-)/abs(G+), are represented by the width of the reaction arrows. The dotted lines represent fluxes that are directionally switched between G- and G+. Quantified metabolites are shown in big characters. Reversible reactions are represented by hexagons and irreversible reactions by diamonds. The direction of the arrows indicates the direction in which the reactions are written in the model. For the sake of clarity, side compounds were omitted. To facilitate interpretation, central carbon metabolism is highlighted in yellow, folate / SAM-SAH cycles in green and nucleotide metabolism in pink. Illustration designed with Cytoscape version 3.9.1
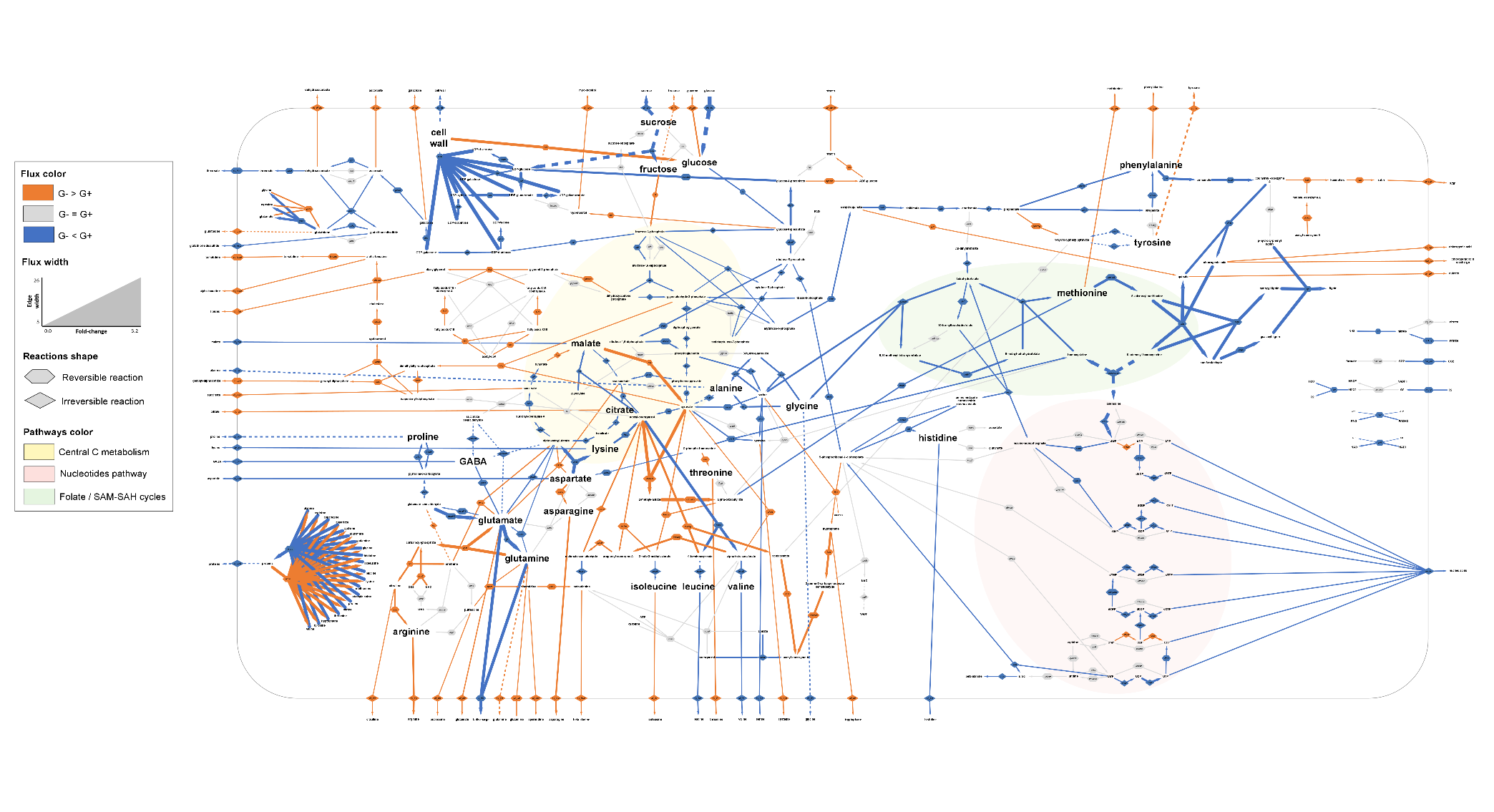

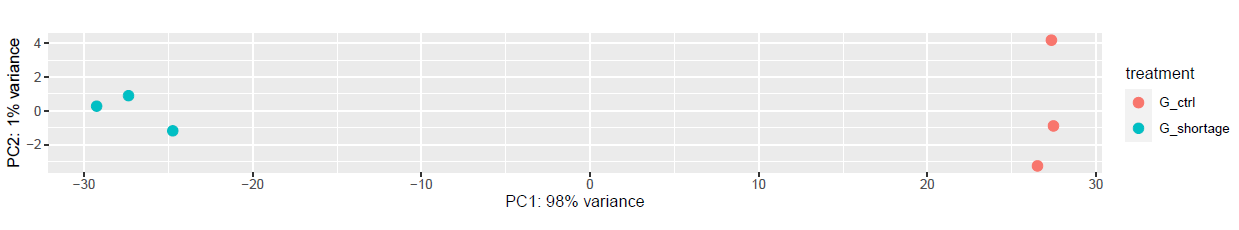

**Supplemental figure S7: Principal Component Analysis (PCA) of RNAseq profile of sequenced samples (n=3) at D6 in G+ and G- conditions. Variance explained by each PC is indicated in brackets.**

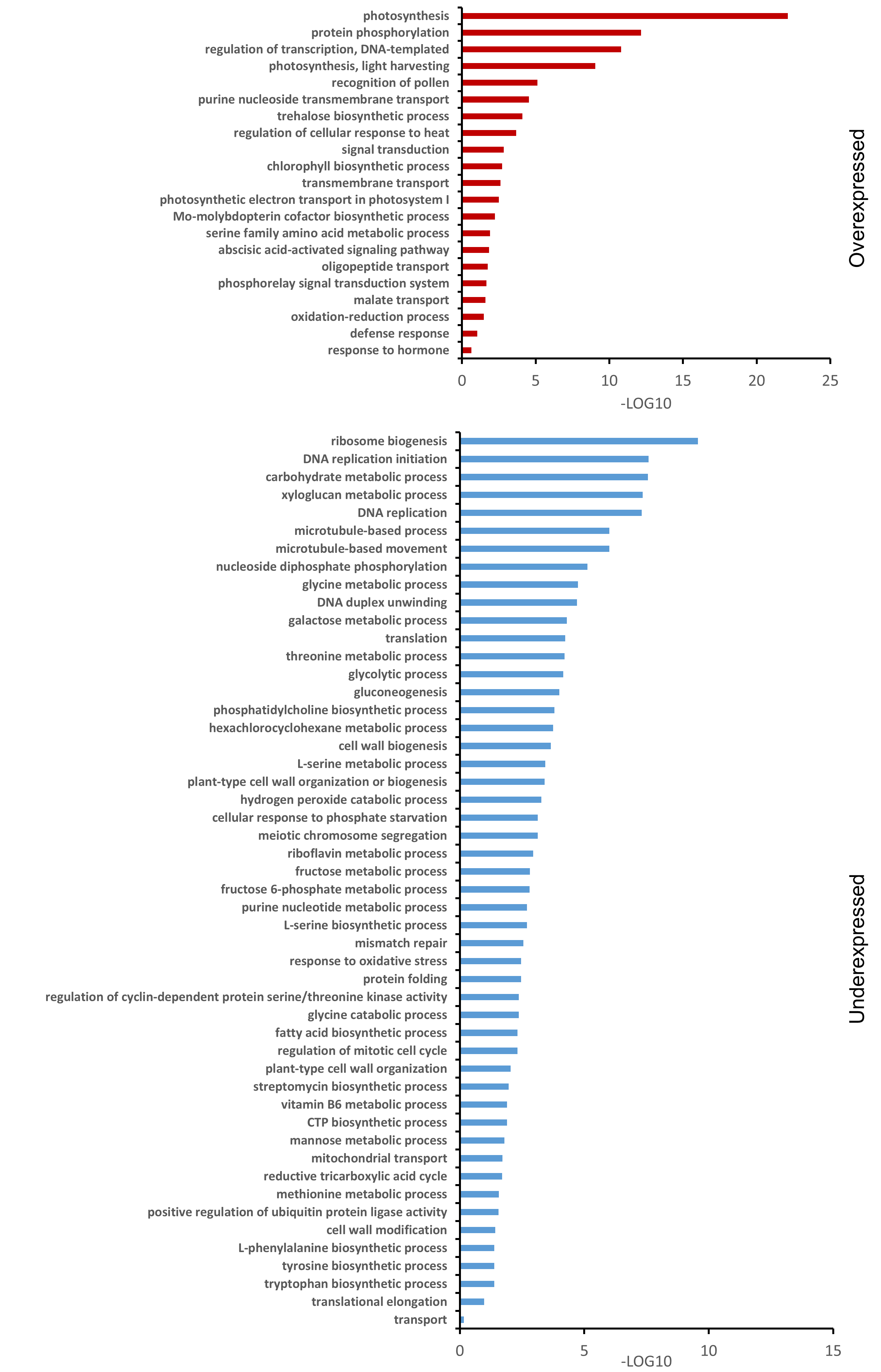

**Figure S8: Gene Ontology (GO) analysis performed on Differentially Expressed Genes (DEGs) identified by comparing G- and G+ conditions. Y-axis represent the biological process GO labels attributed to (A) overexpressed and (B) underexpressed genes. Enrichment values were normalized with the -LOG10 function.**

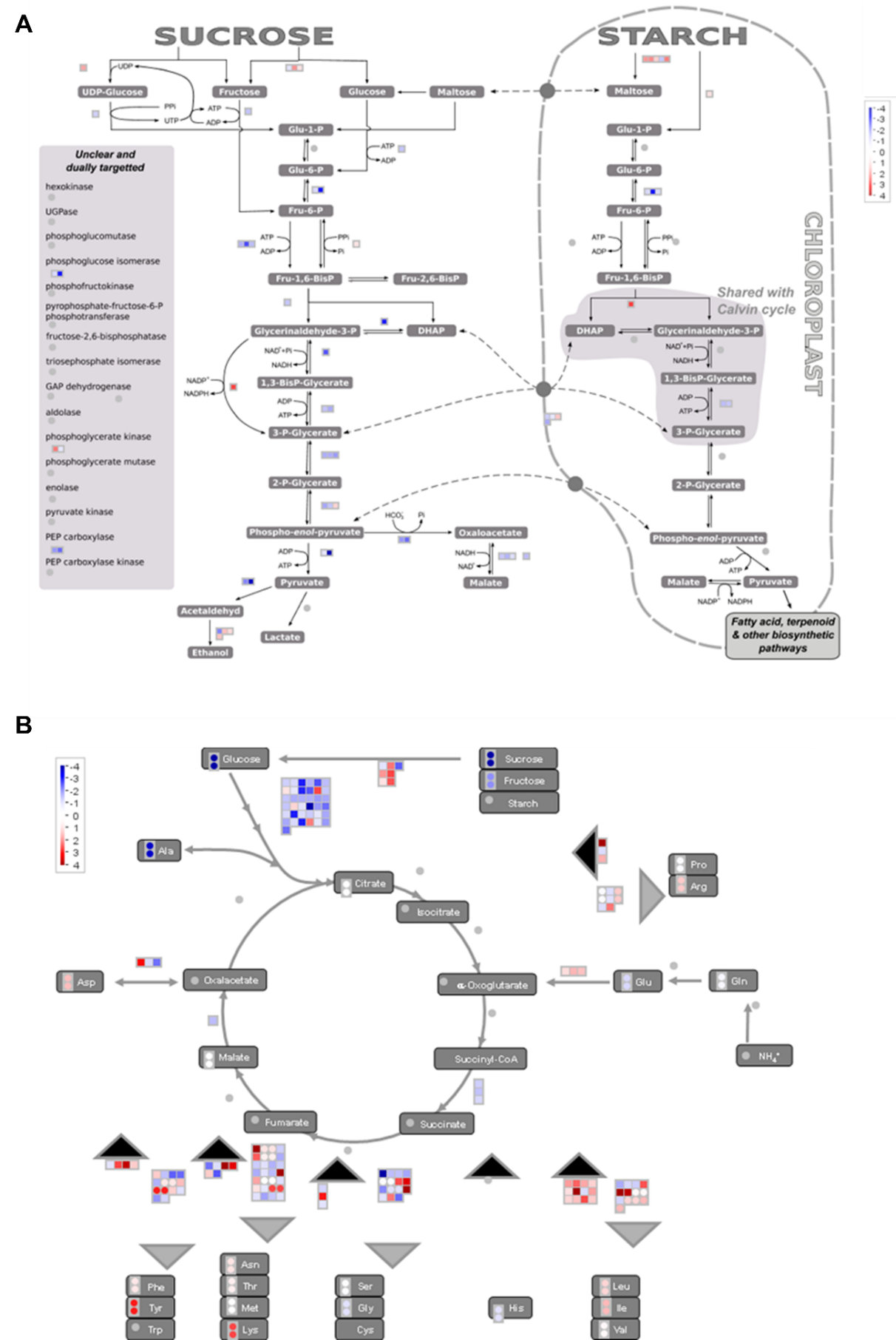

**Figure S9: Mapman view obtain from the DEGs between G- and G+ cells at D6. Visualization focused on glycolysis pathway (A) and TCA cycle (B). Each square represents a gene, each circle represents a metabolite. In red, genes/metabolites showing higher expression/accumulation in G- cells (devprived of sugar); in blue, genes/metabolites showing higher expression/accumulation in G+ cells (control conditions).**

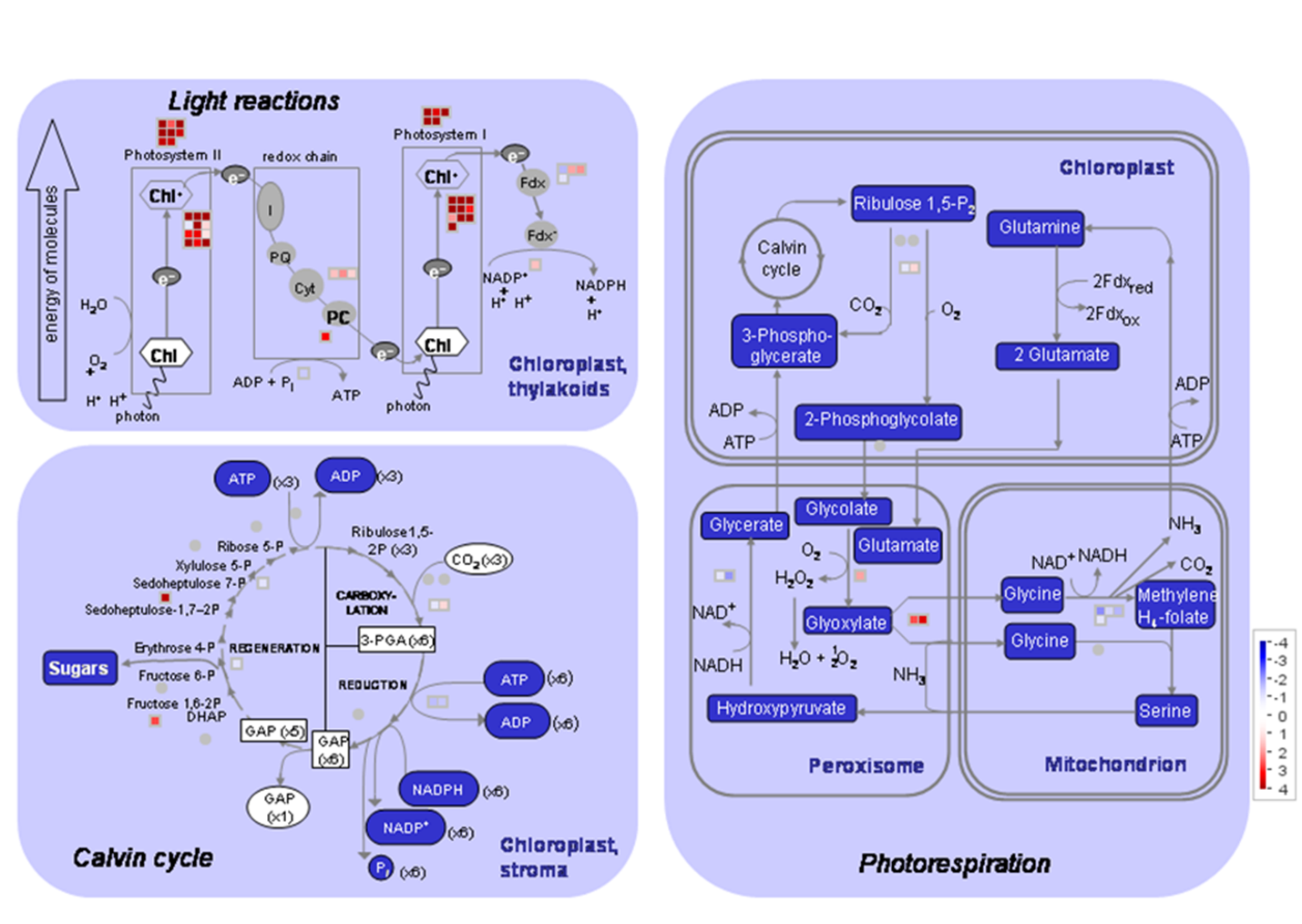

**Figure S10: Mapman representation of energy metabolism overview obtained from the DEGs identified in grapevine cells under carbon limitation conditions compared to control.**

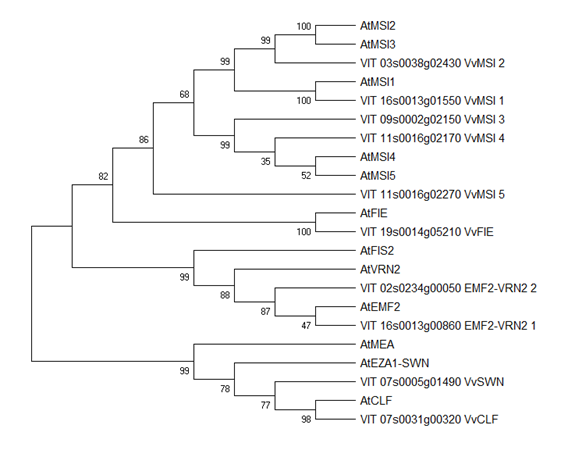

**Figure S11: Phylogenetic tree of PRC2 complex putative homologs identified in Vitis vinifera (Vv) and Arabidopsis thaliana (At). Tree generated with 1,000 bootstrap.**

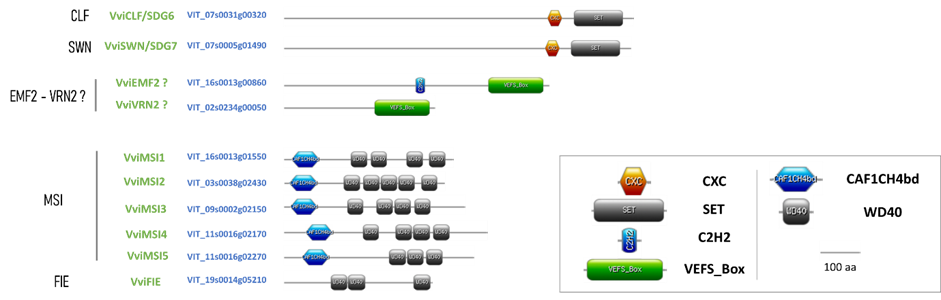

**Figure S12. PRC2 complex putative orthologs identified in grapevine their conserved domains.**

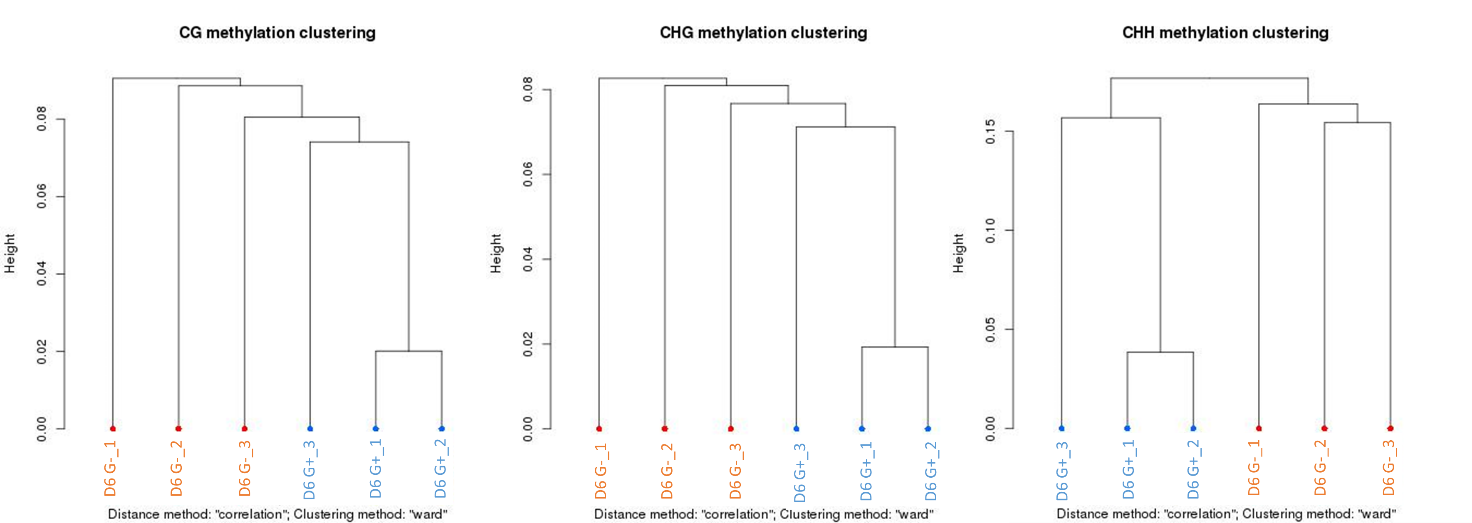

**Figure S13: Dendrogram representation of sample clustering based on their methylome profiles. Methylome analysis performed on G+ (blue) and G- (orange) at D6, clustering using ward method.**
